# A pan-cancer analysis of CpG Island gene regulation reveals extensive plasticity within Polycomb targets

**DOI:** 10.1101/2020.06.04.134858

**Authors:** Yueyuan Zheng, Guowei Huang, Tiago C. Silva, Qian Yang, H Phillip Koeffler, Benjamin P. Berman, De-Chen Lin

**Affiliations:** Department of Medicine, Cedars-Sinai Medical Center, Los Angeles, CA, USA; Department of Pathology, Shantou University Medical College, Shantou, Guangdong, 515041, P.R. China; Center for Bioinformatics and Functional Genomics, Cedars-Sinai Medical Center, Los Angeles, CA, USA; Department of Developmental Biology and Cancer Research, Institute for Medical Research Israel-Canada, Hebrew University-Hadassah Medical School, Jerusalem, Israel

**Keywords:** Polycomb, DNA methylation, gene regulation, cancer, TCGA

## Abstract

CpG Island promoter genes make up more than half of human genes, and a subset regulated by Polycomb-Repressive Complex 2 (PRC2^+^-CGI) become DNA methylated and silenced in cancer. Here, we perform a systematic regulatory analysis of CGI genes across TCGA cancer types, finding that PRC2^+^-CGI genes are frequently prone to transcriptional upregulation as well. These upregulated PRC2^+^-CGI genes control important pathways such as Epithelial-Mesenchymal Transition and TNFα-associated inflammatory response, and have the greatest cancer-type specificity among any class of CGI genes. Using publicly available chromatin datasets and genetic perturbations, we show that transcription factor binding sites (TFBSs) within distal enhancers underlie transcriptional activation of PRC2^+^-CGI genes, in contrast to PRC2-free CGI genes which are predominantly regulated by promoter TFBSs. Surprisingly, a large subset of PRC2^+^-CGI genes that are upregulated in one cancer type are also hypermethylated/silenced in at least one other cancer type, highlighting the high degree of regulatory plasticity likely derived from their complex regulatory patterns during normal development.

## Introduction

Tumorigenesis is a highly complex process driven by both genetic and epigenetic alterations. Among these abnormalities, cancer-specific DNA hypermethylation at CpG-Island (CGI) promoters is perhaps the most well-established epigenetic deregulation. DNA hypermethylation results in transcriptional repression of a large number of genes in cancer. While some are known tumor suppressors, such as *BRCA1*, *MLH1*, and *VHL*, the majority of such hypermethylated genes are “passengers” (little or no functional contribution to cancer biology). In virtually every cancer type, hundreds of CGI promoters are DNA hypermethylated (*1*).

CGI promoters make up a large class of promoters in vertebrate genomes (55-75% of all transcription start sites (TSSs)) (*2*), and only a small fraction are targeted by DNA hypermethylation in cancer. Generally, CGI promoters fall into two major classes: those associated with genes ubiquitously expressed across most cell types (i.e., “housekeeping” genes), and those under complex regulation during embryonic development, which are typically marked with Polycomb group (PcG) proteins (*3*). Both of these classes are unmethylated in embryonic stem cells (ESCs) and most other cell types, but the latter class is prone to DNA hypermethylation in cancer (*4–6*). In fact, PcG-associated genes account for more than 75% of all DNA hypermethylated CGI promoters (*7*). Most of these appear to be passengers, albeit there is a subset with tumor suppressor function, including *SFRP5 (8)*, *GATA5 (9)*, and *RUNX3 (10)*. A subgroup of highly regulated developmental transcription factors (TFs) have much longer (> 5 kilobase) regions of de-methylated and CGI-containing DNA (*11*), and these “DNA methylation valleys” (DMVs) also gain methylation in cancer (*12, 13*).

Initially discovered in Drosophila melanogaster, PcG factors play a major role in the regulation of cell fate and differentiation (*14*). They form multiple Polycomb Repressive Complexes (PRCs), including PRC1 and PRC2. Compared with PRC1, the function and regulation of PRC2 is more extensively characterized and better understood (*14*). In mammals, PRC2 is ubiquitously expressed and preferentially binds to CGI promoters to mediate mono-, di- and tri-methylation of histone H3 lysine 27 (H3K27me1/me2/me3) (*14, 15*). Among them, H3K27me3 is considered as a hallmark of PcG-dependent transcriptional repression. The methyltransferase activity of PRC2 is regulated by three core components, enhancer of zeste homologue 1 (EZH1) or EZH2, suppressor of zeste 12 (SUZ12) and embryonic ectoderm development (EED) (*15*). Mechanistically, the susceptibility of PcG-occupied CGI promoters to DNA hypermethylation may be related to the capability of EZH2 in recruiting DNMT3a (*16*). Functionally, in addition to maintaining transcriptional repression, PRC2 also establishes a unique “bivalent” chromatin in many unmethylated CGI promoters, which is especially prominent in ESC (*17*). Harboring both H3K27me3 and active histone marks (H3K4me2/3), bivalent chromatin is considered to maintain a low but poised transcriptional state either for rapid activation in specific developmental context or long-term repression in other cell types.

While numerous studies have focused on the hypermethylation and epigenetic silencing of PRC2 associated genes, very few have looked systematically at how the entire class of PRC2-occupied CGI promoters are dysregulated in cancer. Isolated studies have indicated that these promoters might not only be prone to silencing, but also to transcriptional activation in cancer (*18*). For example, in colon cancer, many stem cell regulators and proliferation-promoting factors with bivalent promoters became active after losing PcG mark H3K27me3 (*19*). In addition, a few PRC2-regulated genes, such as the well-defined leukemic oncogene *HOXA9*, have been observed to be either DNA hypermethylated or transcriptionally upregulated in different cancer types (*20*). DLX5, which contains a bivalent promoter in ESCs and most normal tissues, is converted to an active chromatin state in squamous cancers and can promote proliferation and migration in these cancer types (Manuscript by same authors, under revision in this Journal). These observations indicate that although Polycomb-occupied promoters are well-known to become DNA hypermethylated and epigenetically silenced in cancer, the opposite change may also be common in cancer and play a role in tumor biology. However, the prevalence of this type of transcriptional activation (i.e., how many promoters are affected) in the full spectrum of human cancers, the biological significance of the alteration, and the underlying mechanisms of the transcriptional activation are unknown, possibly because this change has little or no effect on promoter DNA methylation.

To address these questions, using tumor and nonmalignant samples across pan-cancer types, we analyzed comprehensively the transcriptomic and epigenomic characteristics of PRC2-occupied CGI (referred to as PRC2^+^-CGI) genes and PRC2-free CGI (referred to as PRC2^−^-CGI) genes. In addition to the expected epigenetically silenced genes, we also found a significant subset that was upregulated in cancer, and investigated the tissue-specificity and functional significance of this new class of genes, as well as the regulatory mechanisms underlying their activation in the context of cancer biology.

## Results

### Systematic identification of transcriptionally deregulated PRC2^+^- and PRC2^−^-CGI genes across human cancers

To characterize alterations in PRC2-occupied CGI (referred to as PRC2^+^-CGI) promoters in cancer, we first curated a comprehensive set of TSSs consisting of 101,819 loci from GENCODE and 43,164 from the FANTOM4 Cap-Assisted Gene Expression dataset (**Fig. S1**). A total of 53,860 promoters associated with these TSSs were covered by methylation probes on the HM450K methylation array, and approximately 70% (35,686/53,860) of these promoters overlapped a CGI region. Among CGI promoters, more than 20% (7,573/35,686) were defined as PRC2^+^ in ESCs, based on the annotation of repressed/bivalent ChromHMM states (*21*) as well as EZH2/SUZ12-binding (**Fig. 1A**). We analyzed the same datasets and identified 21,226 CGI promoters without PRC2-occupancy (termed PRC2^−^-CGI, **Fig. 1A** and **Materials and Methods**).

**Fig. 1.**
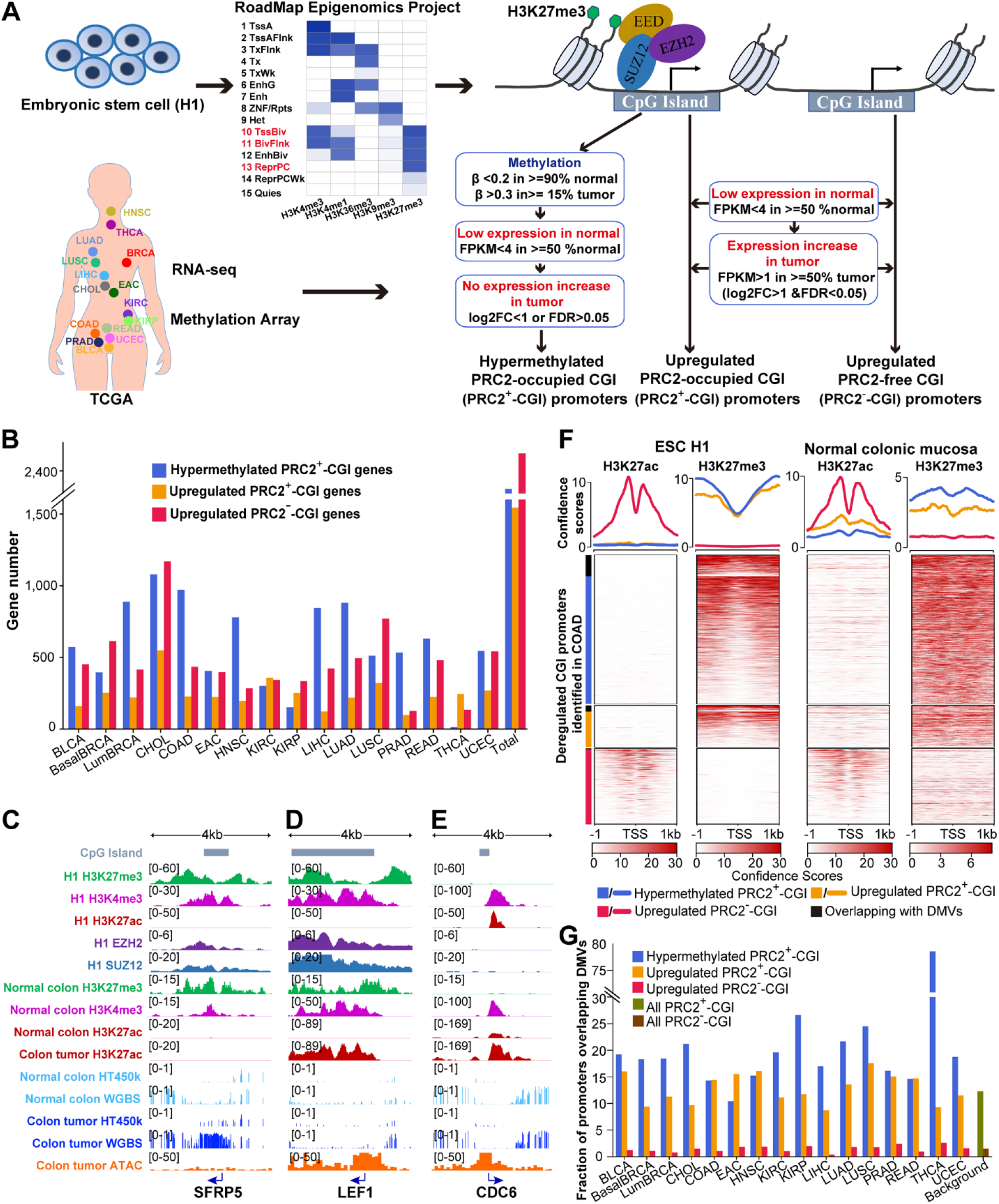
Systematic identification of transcriptionally deregulated PRC2^+^-CGI and PRC2^−^-CGI genes across human cancers. **(A)** An integrated pipeline for identification of three different classes of CGI promoters. **(B)** The numbers of each class of CGI genes in each TCGA cancer type. **(C-E)** Integrative Genomics Viewer (IGV) plots show representative colon adenocarcinoma (COAD) genes for three classes: **(C)** hypermethylated PRC2^+^-CGI, **(D)** upregulated PRC2^+^-CGI and **(E)** upregulated PRC2^−^-CGI. ChIP-Seq data are from Roadmap and ENCODE projects. Methylation data are from TCGA. **(F)** Three classes of genes identified in COAD show similar H3K27ac and H3K27me3 patterns between ESC and normal colonic mucosa. Promoters overlapping with ESC DNA Methylation Valleys (DMVs) from Xie *et.al (12)* are marked with black bars. **(G)** Fraction of all promoters overlapping DMV regions.

Using ESC chromatin marks to define PRC2^+^ and PRC2^−^ gene classes is a common strategy, given that H3K27me3 ChIP-Seq becomes more diffuse in differentiated cell types and PRC2 ChIP-Seq has been attempted in very few differentiated cell types (*6, 7, 22*). Our approach is well-justified, since most regions that are PRC2-occupied in ESCs retain H3K27me3 in most differentiated cell types (*12*). Nevertheless, we sought to confirm that these annotations were representative of the PRC2-occupied state in normal tissues. Since PRC2-occupancy is associated with repressed/bivalent transcription, we reasoned that PRC2^+^-CGI genes should have minimal expression in normal tissues. We thus analyzed the mRNA levels of PRC2^+^-CGI genes in TCGA nonmalignant samples with available histone markers from the NIH Roadmap project (including colonic mucosa, lung, breast epithelium, rectum, esophagus, uterus and liver). As anticipated, H3K27ac signals of PRC2^+^-CGI genes were significantly correlated with their mRNA expression levels, and most were low for both. An FPKM value of 4 prominently separated PRC2^+^-CGI genes with distinct H3K27ac levels (**Fig. S2A**), and the H3K27me3 mark, a hallmark of PRC2-occupancy, was only positive in PRC2^+^-CGI genes with FPKM < 4 (**Fig. S2B**). As expected, only an average of 22.4% PRC2^+^-CGI genes had FPKM > 4 in nonmalignant tissues. These results are consistent with previous reports that a large fraction of silenced H3K27me3-covered regions are shared between ESCs and differentiated cells (*12*).

Upon cataloging the class of PRC2^+^-CGI genes (n=4,378, covering 7,573 promoters) in normal tissues, we next investigated the cancer-associated changes in these genes by analyzing the transcriptomic data and DNA methylation data of TCGA cancer types (**Fig. 1A** and **Materials and Methods**). A total of 16 TCGA cancer types with sufficient nonmalignant samples (n >= 5) were used, including bladder cancer (BLCA), basal breast cancer (BasalBRCA), luminal breast cancer (LumBRCA), cholangiocarcinoma (CHOL), colon adenocarcinoma (COAD), esophageal adenocarcinoma (EAC), head and neck squamous cell carcinoma (HNSC), kidney renal clear cell carcinoma (KIRC), kidney renal papillary cell carcinoma (KIRP), liver hepatocellular carcinoma (LIHC), lung adenocarcinoma (LUAD), lung squamous cell carcinoma (LUSC), prostate adenocarcinoma (PRAD), rectum adenocarcinoma (READ), thyroid carcinoma (THCA) and uterine corpus endometrial carcinoma (UCEC) (**Table S1**).

We first identified hypermethylated PRC2^+^-CGI promoters (**Fig. 1B** and **Table S2**) using criteria based on those developed by the TCGA consortium (*23*). Consistent with well-established findings (*1*), almost 52% of PRC2^+^-CGI genes (2,274 of 4,378) became hypermethylated in at least one cancer type, corresponding to 4,260 promoters (60% of all PRC2^+^-CGI promoters). Most cancer types (12/16) had > 400 hypermethylated PRC2^+^-CGI genes (**Fig. 1B**), confirming the pervasiveness of this type of epigenetic silencing of PRC2^+^-CGI promoters across human cancers. For the example of COAD, we verified known hypermethylated tumor suppressors (**Table S3A**), such as *SFRP5 (8)*, *GATA5 (9)*, and *RUNX3 (10)*. As shown in **Fig. 1C**, *SFRP5* harbors a bivalent promoter in both H1 cells and normal colon tissue, with both H3K27me3 and H3K4me3 signals, but devoid of H3K27ac. In colon cancer, this promoter becomes DNA hypermethylated and inaccessible (undetectable ATAC-Seq signal).

In addition to the epigenetically silenced genes, more than 35% of PRC2^+^-CGI genes (1,543 of 4,378, associated with 2,891 promoters) were upregulated across cancer types (**Fig. 1B**). On average, each cancer type possessed 245 (ranging from 98 to 549) such upregulated PRC2^+^-CGI genes (**Table S2**). Consistent with a previous study (*19*), our analyses in colon cancer revealed many upregulated PRC2^+^-CGI genes associated with WNT signaling pathway (**Table S3B**), such as *LEF1*, *LGR5*, *WNT2*. Shown as an example, the *LEF1* gene is marked simultaneously with H3K27me3 and H3K4me3 but without H3K27ac in both ESCs and normal colon tissue. In contrast, it is transcriptionally active in colon tumors and is accompanied by high accessibility, conspicuous H3K27ac level and low DNA methylation (**Fig. 1D**).

We next sought to characterize the PRC2^−^-CGI gene class, which is considered to have relatively ubiquitous expression across different cell types (*3*). We applied the identical criteria of differential expression that were used for PRC2^+^-CGI genes and identified ~2,500 upregulated PRC2^−^-CGI genes across cancers (**Table S2**). Most cancer types (10/16) had > 400 such upregulated PRC2^−^-CGI genes (**Fig. 1B**). For example, *CDC6*, an essential factor for DNA replication, harbors an active promoter marked with both H3K27ac and H3K4me3 in both ESCs and normal tissue, and becomes transcriptionally upregulated in 75% (12/16) cancer types (**Table S3C**), such as colon cancer (**Fig. 1E**).

We further confirmed the consistency in chromatin state between ESCs and normal tissues in the three classes of CGI genes. As anticipated, the H3K27me3 signals in ESCs were high in PRC2^+^-CGI promoters but undetectable in PRC2^−^-CGI promoters, and the opposite pattern was observed in the distribution of H3K27ac signals (**Fig. 1F**). Importantly, a highly similar profile of both H3K27me3 and H3K27ac signals was seen in almost all normal tissues (**Fig. 1F** and **Fig. S3**). This result confirms that the promoter classes investigated in this study largely maintain their ESC chromatin states during normal development.

As described above, DNA methylation valleys (DMVs) represent a functionally important subgroup of PRC2^+^-CGI genes that have hypermethylated promoters in cancer. Looking specifically at DMVs, we found that they were associated both with hypermethylation and upregulation of PRC2^+^-CGI genes in similar ratios (**Fig. 1F-G**), although hypermethylation tended to be slightly more enriched. Overall, DMVs were represented at comparable proportions in these classes as they were in PRC2^+^ genes overall (green bar in **Fig. 1G**). Given the functional importance of these genes during development, upregulated PRC2^+^ DMV genes may represent a small but important class of cancer-promoting genes.

### Upregulated PRC2^+^-CGI genes have increased promoter H3K27ac and accessible chromatin in cancer

We next identified chromatin changes at CGI promoters in cancer, using DNA methylation data and ATAC-Seq data from TCGA, as well as H3K27ac ChIP-Seq from individual studies. ATAC-Seq and H3K27ac are particularly useful, as they are both strongly correlated with transcriptional activity of cis-regulatory elements. The TCGA ATAC-Seq project (*24*) did not include non-malignant tissues for comparison, but in tumors we could clearly see that the hypermethylated PRC2^+^-CGI promoters were inaccessible, whereas both the upregulated PRC2^+^-CGI and PRC2^−^-CGI promoters were significantly accessible (**Fig. 2A**). By re-analyzing data from 4 nonmalignant colonic crypts and 17 primary colon cancer cells (GSE77737), we were able to measure the cancer-specific changes in H3K27ac for the three classes of genes (**Fig. 2B** and **Fig. S4A**). Hypermethylated PRC2^+^-CGI promoters had undetectable levels of H3K27ac in both nonmalignant and tumor samples, whereas most upregulated PRC2^+^-CGI promoters had low H3K27ac in nonmalignant samples and a gain of the mark in tumors. The upregulated PRC2^−^-CGI promoters also gained H3K27ac in tumors, but were typically already high for the mark in the nonmalignant samples. In another cohort of 7 KIRC tumors with matched adjacent nonmalignant tissues (GSE86095), similar patterns were observed across the three gene classes (**Fig. 2C** and **Fig. S4B**).

**Fig. 2.**
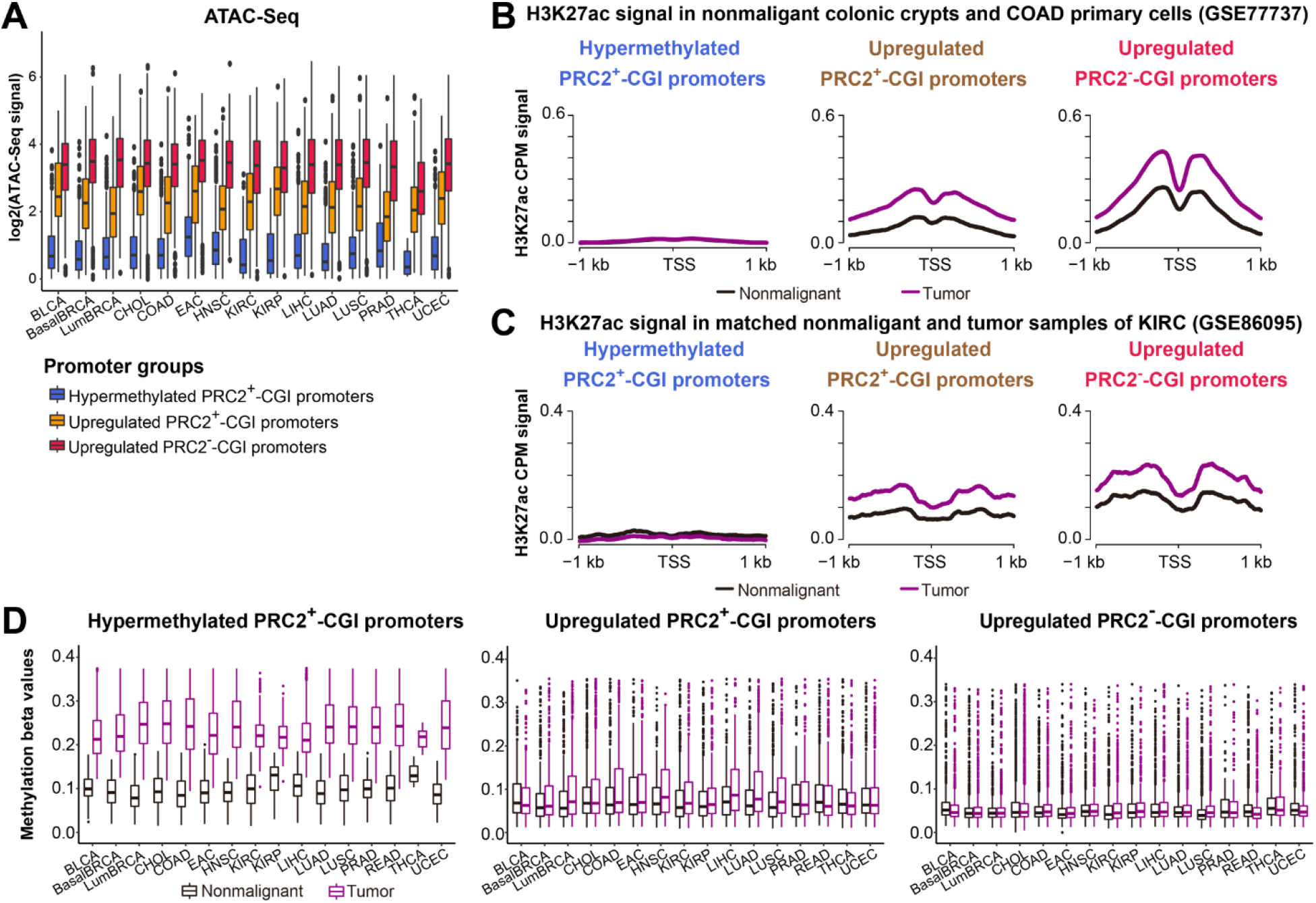
Upregulated PRC2^+^-CGI genes have increased promoter H3K27ac levels and accessible chromatin in tumors. **(A)** TCGA ATAC-Seq signals for each of the three CGI promoter classes in each cancer type. **(B)** Aggregation plots of averaged non-malignant and primary colon cancer cells H3K27ac ChIP-Seq signals (GSE77737) for each of the three CGI promoter classes from COAD (see **Fig. S4** for details). **(C)** Similar line plots generated using H3K27ac ChIP-Seq signal from matched non-malignant and tumor pairs of KIRC (GSE86095, see **Fig. S4** for details). **(D)** Methylation beta values for each of the three CGI promoter classes in nonmalignant and tumor samples across TCGA cancer types. The top 5% of outliers in each group were outside this range and not plotted.

At the DNA methylation level, as anticipated, all three classes of CGI promoters had low methylation across nonmalignant tissues (**Fig. 2D**). The increased DNA methylation at PRC2^+^-CGI hypermethylated promoters was evident in tumors, while methylation levels were largely unchanged in upregulated PRC2^+^- and PRC2^−^-CGI promoters. Interestingly, methylation levels were slightly higher in upregulated PRC2^+^-CGI promoters in several cancer types. This direction of change goes counter to the usual anti-correlation between DNA methylation and expression, but is consistent with observations in another study analyzing TCGA data (*18*).

### Upregulated PRC2^+^-CGI genes have the highest levels of cancer-type specificity and regulatory plasticity

We next sought to compare expression levels for the three CGI gene classes to determine their specificity with respect to cancer type. Consistent with earlier reports (*7*), hypermethylated PRC2^+^-CGI genes were slightly downregulated in TCGA tumors relative to adjacent nonmalignant tissues (**Fig. 3A-B** and **Fig. S5A-B**). While both upregulated PRC2^+^-CGI and PRC2^−^-CGI classes were pre-selected to have an expression increase of at least 2-fold, the PRC2^+^-CGI set showed higher relative increases in 11/15 cancer types (**Fig. 3A**). For example in COAD, ~40% (92/228) of upregulated PRC2^+^-CGI genes were increased by more than 4-fold; in comparison, only 15% (62/424) of PRC2^−^-CGI class showed a 4-fold increase (**Fig. 3B**). The higher induction of a subset of PRC2^+^-CGI implicates its biological significance in cancer, with their expression levels potentially being under positive selection.

**Fig. 3.**
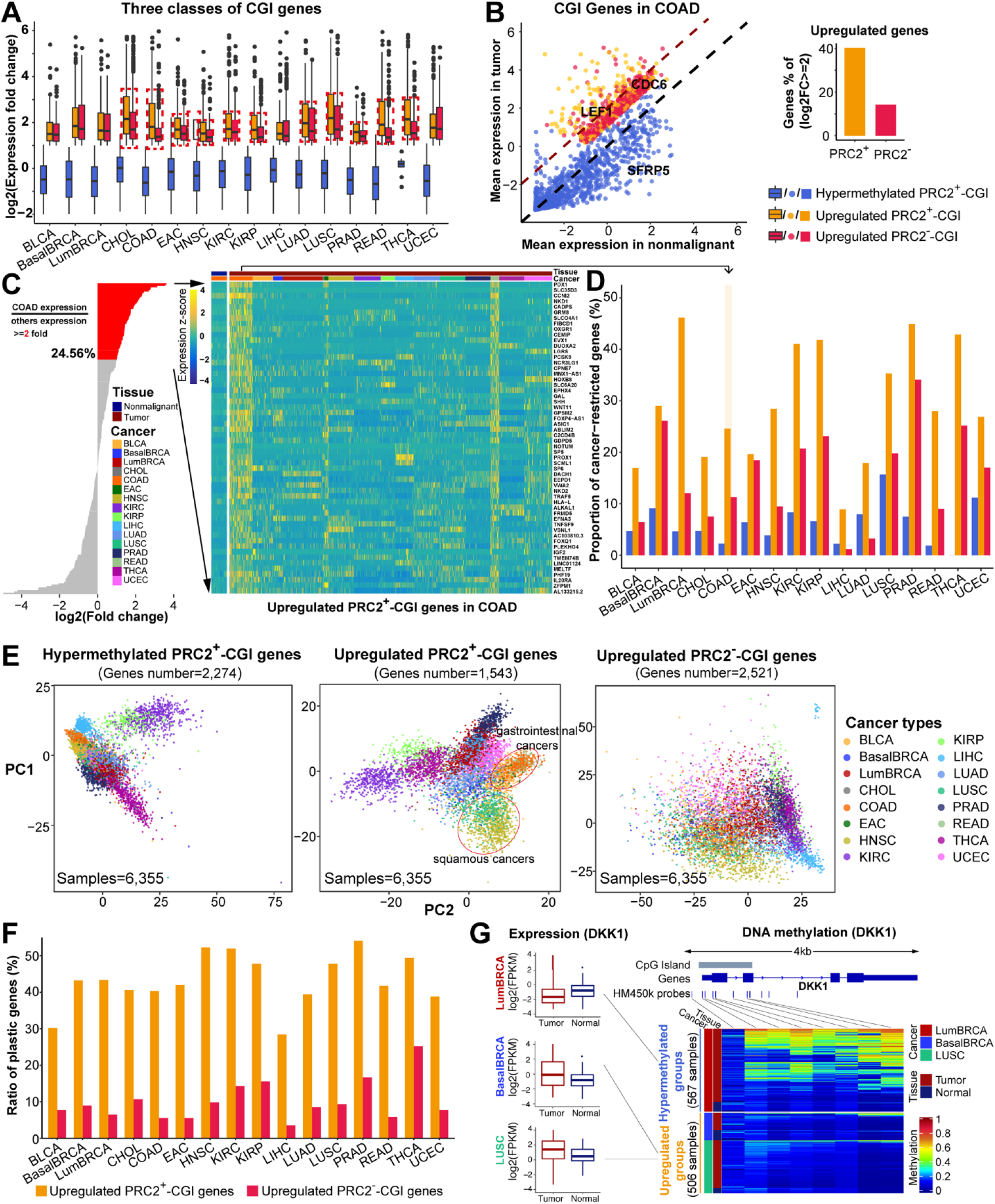
Upregulated PRC2^+^-CGI genes have the highest degrees of cancer-type specificity and regulatory plasticity. **(A)** Expression fold change between tumor and nonmalignant samples, stratified by CGI promoter classes. Red boxes highlight the 11 cancer types where PRC2^+^-CGI genes have the highest overexpression. **(B)** Individual genes plotted for COAD, exemplary genes from **Fig. 1C-E** were highlighted. **(C)** Cancer-type-restricted genes are identified based on expression fold change between a specific cancer type (COAD in this example) versus all other cancer types. Fold-change is shown on the left, with all genes sorted by fold-change and those with >=2 (“cancer-type-restricted genes”) shown in red. The 56 cancer-type-restricted genes for COAD are shown as a heatmap on the right. **(D)** The percentage of cancer-type-restricted genes from each gene class, shown by cancer type. **(E)** Plastic genes were defined as those assigned to the upregulated group in one cancer type, and hypermethylated/not upregulated in another. The percentage of plastic genes is stratified by cancer type where upregulation occurs. **(F)** TCGA methylation and expression data are shown for a plastic gene (*DKK1*) that is upregulated in BasalBRCA and LUSC, and hypermethylated/downregulated in LumBRCA. **(G)** PCA analyses using expression values from each of the three CGI gene classes. Cancer types clustered by PRC2^+^-CGI genes are highlighted by circles, such as squamous cancers (i.e., LUSC, HNSC and a subset of BLCA) and gastrointestinal cancers (i.e., EAC, COAD and READ).

To investigate cancer-type specificity, we determined the extent and significance of expression differences between each cancer type vs. all others, and labeled those with fold-change >=2 as cancer-type-restricted genes (**Fig. 3C**). DNA hypermethylated genes are known to have some cancer-type specificity but its degree is relatively low (*23*), and indeed this class had the lowest percentage of cancer-type-restricted genes in all but 2 cancer types (**Fig. 3D**). Upregulated PRC2^+^-CGI genes had the largest fraction of cancer-type-restricted genes across all cancer types, on average 2.67 fold higher than that of PRC2^−^-CGI genes.

Interestingly, nearly half (762/1,543) of upregulated PRC2^+^-CGI genes were assigned to the hypermethylated and non-upregulated class in another cancer type. In contrast, only 8.5% (300/2,521) of upregulated PRC2^−^-CGI genes showed this type of regulatory plasticity. Indeed, in every cancer type, the fraction of “plastic genes” was much higher in PRC2^+^- than PRC2^−^-CGI class (**Fig. 3E**). As an example, the PRC2^+^-CGI promoter of *DKK1* is hypermethylated and silenced in LumBRCA, but remains unmethylated and is upregulated in both BasalBRCA and LUSC (**Fig. 3F**), consistent with earlier reports in ER-/PR-negative breast cancer (*25*), lung cancer (*26*), and LumBRCA cancer (*27*). Additional examples of plastic genes are shown in **Fig. S5C-D**.

We next performed unsupervised clustering of all 6,355 TCGA tumor samples with PCA analysis using expression values for each of the three gene classes independently (**Fig. 3G**). Cancer types were separated most clearly based on the upregulated PRC2^+^-CGI class, as predicted by the higher percentage of cancer-type-restricted genes in this class. A similar result was obtained by PCA analysis of ATAC-Seq data (**Fig. S5E**). Moreover, the PCA analysis of PRC2^+^-CGI genes revealed additional patterns which were not found by clusterings based on the other two classes (**Fig. 3G**): i) cancer subtypes derived from the same tissues were split into different molecular clusters, such as breast (BasalBRCA and LumBRCA), lung (LUAD and LUSC) and kidney cancer (KIRC and KIRP); ii) cancer types sharing either similar cell-of-origins or lineages were clustered together across different anatomical locations, such as squamous cell carcinoma (including LUSC, HNSC and a subset of squamous-like BLCA) and gastrointestinal cancers (including EAC, COAD and READ).

### Upregulated PRC2^+^-CGI and PRC2^−^-CGI genes control distinct sets of biological pathways in cancer

We explored differential biological functions of upregulated PRC2^+^-CGI and PRC2^−^-CGI genes in cancer using Hallmark pathway enrichment by setting one of the two gene classes as the foreground and the other as the background. This revealed distinct sets of biological pathways enriched in the two classes of genes (**Fig. 4A-B**). Among the top-ranked PRC2^+^-CGI-enriched pathways across multiple cancer types, we found “Epithelial mesenchymal transition (EMT)”, “KRAS signaling up”, and “TNFα signaling via NF-KB” pathways, while other pathways such as “Estrogen response early” were highly cancer-type specific (**Fig. 4A**). In contrast, PRC2^−^-CGI genes had little cancer-type specificity (**Fig. 4B**), and the three top ranked pathways were all cell-cycle related which were enriched across 15/16 cancer types (“E2F targets”, “G2M checkpoint” and “Mitotic spindle”, **Fig. 4B**). This marked difference in cancer-type specificity was even more apparent at the individual gene level, with 50%~61.2% of PRC2^−^-CGI genes enriched in the top three pathways shared by over half of all cancer types, compared to only 2.1%-8.1% of genes enriched in the top three PRC2^+^-CGI pathways (**Fig. S6**). These findings highlight that upregulated PRC2^+^-CGI genes control distinct sets of biological pathways in a cancer-type specific manner, consistent with their high cancer-type expression specificity described above.

**Fig. 4.**
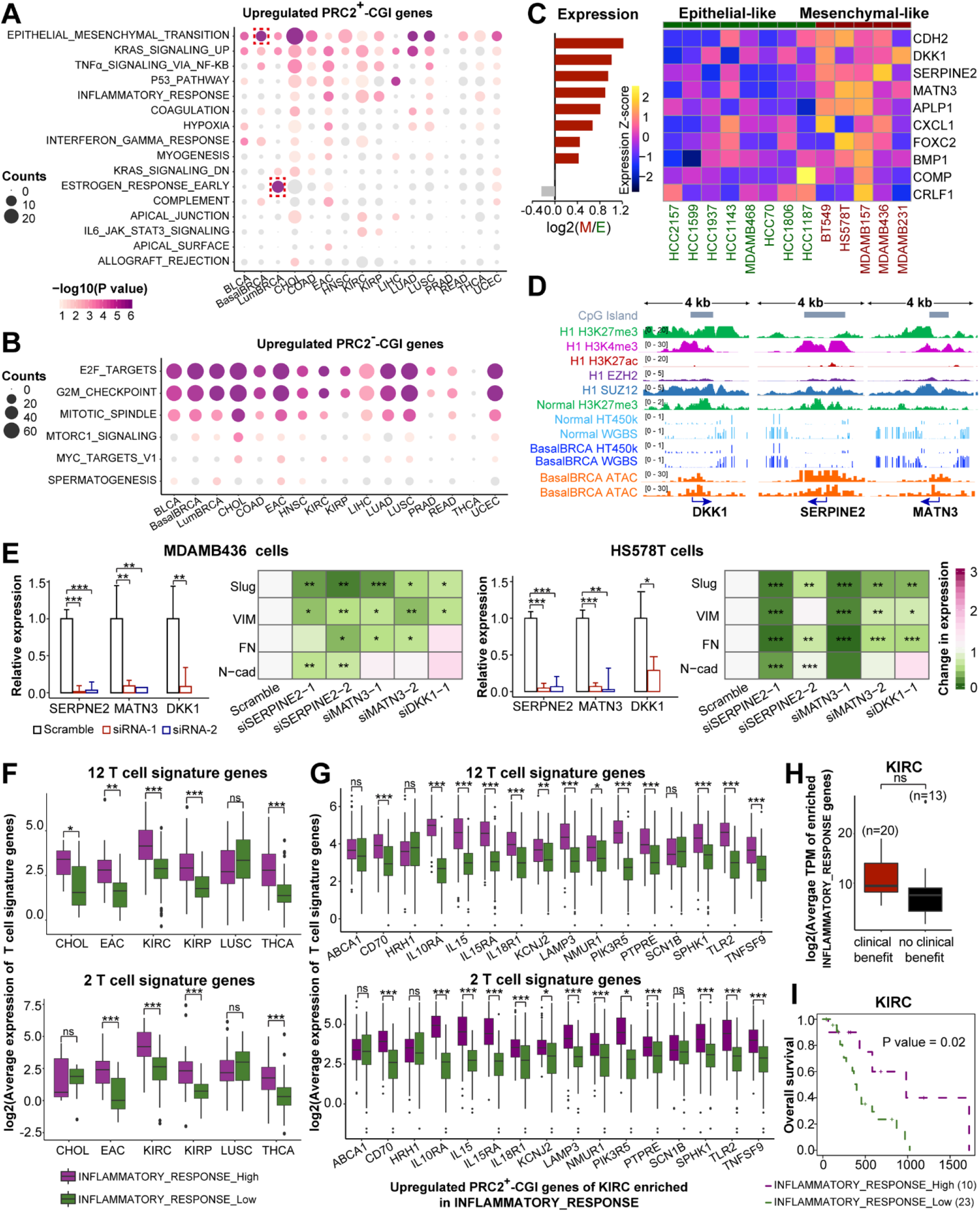
Upregulated PRC2^+^- and PRC2^−^-CGI genes control distinct sets of biological pathways in cancer. **(A-B)** Hallmark pathway enrichment results for **(A)** upregulated PRC2^+^-CGI genes and **(B)** upregulated PRC2^−^-CGI genes. “EMT” and “Estrogen response early” are highlighted in the two breast cancer subtypes. **(C)** Expression of the 10 BasalBRCA upregulated PRC2^+^-CGI genes in the EMT pathway are shown in 13 BasalBRCA cell lines annotated by an established consensus EMT classification (see text). **(D)** IGV plots for the top 3 novel EMT genes from panel **C**, showing PRC2 marks in ESCs and chromatin marks in normal breast epithelium and BasalBRCA tumors. DNA methylation is averaged over all TCGA samples. **(E)** siRNA loss-of-function assays for the top three novel genes followed by expression measurement of established mesenchymal markers. Data are presented as mean±SD of two replicates (left), showing p values for each individual gene and replicate, as determined by t-test. p<0.001, ***; p<0.01, **; p<0.05, *. **(F-G)** Average expression of CD8+ T-cell signature genes in the six cancer types enriched in the inflammatory response pathway from panel **A**, showing the top and bottom 20% of tumors based on either the average **(F)** or the individual gene-level **(G)** expression of upregulated PRC2^+^-CGI genes in the inflammatory response pathway. p values were determined by t-test. p<0.001, ***; p<0.01, **; p<0.05, *; p>0.05, ns. Expression datasets are obtained from Miao *et.al. (35)*. **(H)** The average expression of upregulated PRC2^+^-CGI genes in the inflammatory response pathway in KIRC patients with differential response to immune checkpoint therapies. **(I)** Kaplan-Meier survival plot analyzing the average expression of upregulated PRC2^+^-CGI genes in inflammatory response pathway using the same cohort of KIRC patients.

We next sought to functionally validate the pathway enrichment results of PRC2^+^-CGI genes, using the EMT pathway in BasalBRCA as an example. We chose this example since it was the most significantly enriched pathway across multiple cancer types, and because EMT has well-defined biological significance in BasalBRCA, which also provides multiple cell line models for experimental interrogation *in vitro* (*28*). An established consensus classification of EMT based on expression data (*29*) was used to identify epithelial- and mesenchymal-like basal breast cell lines, and we selected the 8 mesenchymal- and 5 epithelial-like cell lines that were also profiled by the Cancer Cell Line Encyclopedia (**Fig. S7A**). Of the 10 PRC2^+^-CGI upregulated genes that were identified in the enriched EMT pathway in BasalBRCA, we found that 8 had higher expression in the mesenchymal-compared to the epithelial-like lines (**Fig. 4C**), including several known EMT-promoting factors, including CDH2, CXCL1, FOXC2 and BMP1 (28, 30, 31). Most of these 10 genes showed clear PRC2 occupancy in ESCs and normal breast tissue, as well as high chromatin accessibility in BasalBRCA TCGA tumors (**Fig. 4D** and **Fig. S7B**).

Three of the top four genes enriched in PRC2^+^-CGI EMT pathway had not been functionally implicated in EMT phenotype in breast cancer (*SERPINE2*, *DKK1*, *MATN3*). We performed siRNA loss-of-function assays for these genes in two mesenchymal-like cell lines (MDAMB436 and HS578T) that had high endogenous levels of these three factors (**Fig. 4C**). In both cell lines, knockdown of either SERPINE2, DKK1 or MATN3 by individual siRNAs markedly reduced the expression of known mesenchymal markers (**Fig. 4E**). These results validate the biological contribution of PRC2^+^-CGI genes to the EMT pathway and identify three novel PRC2^+^-CGI factors (SERPINE2, DKK1, MATN3) with EMT-promoting function in basal breast cancer. As described above, *DKK1* is also notable as a plastic gene and becomes hypermethylated/silenced in LumBRCA (**Fig. 3F**).

In addition to the EMT pathway, we noted that two immune-related pathways were ranked among top 5 in the PRC2^+^-CGI class, namely “TNFα signaling via NF-KB” and “Inflammatory response”. Since both pathways have well-defined roles in anti-tumor immunity and contribute to immune-checkpoint blockade therapy (*32*), this result raises the possibility that the activation of these two pathways by PRC2^+^-CGI genes might be associated with increased immunity against cancer cells. We thus analyzed the cytotoxic activity of infiltrating CD8+ T cells based on two independent, well-established gene signatures (*33, 34*) in the 6 cancer types enriched for “Inflammatory response” (from **Fig. 4A**). Tumor samples with higher average expression of “Inflammatory response” PRC2^+^-CGI genes showed higher cytotoxic activity of intratumoral CD8+ T cells in most cancer types (**Fig. 4F**), and this was the case for most of the 16 “Inflammatory response” genes individually (**Fig. 4G**). The same was true for PRC2^+^-CGI genes of the “TNFα signaling via NF-KB” pathway (**Fig. S7C-D**), albeit these two pathways share a number of genes in common. We next explored whether activation of these pathways by PRC2^+^-CGI genes was associated with the response to immune-checkpoint blockade therapy. Of all the enriched cancer types, only KIRC patients had available RNA-Seq data prior to immuno-therapy (*35*). Compared with patients with no clinical benefit from anti-PD-1 therapy, those showing clinical response expressed higher “Inflammatory response” PRC2^+^-CGI genes albeit without reaching statistical significance (**Fig. 4H**), and patients with higher expression of these genes also had better overall survival following anti-PD-1 therapy (**Fig. 4I**). A similar trend of TNFα pathway was also observed (**Fig. S7E-F**).

### Upregulated PRC2^+^-CGI genes are linked to distal enhancers targeted by specific transcription factor binding sites (TFBSs)

We next identified candidate upstream regulators of PRC2^+^-CGI v.s. PRC2^−^-CGI genes using TFBS motif enrichment analysis. For promoter TFBSs, we used the promoter regions as described above. Enhancer elements associated with a particular promoter can act over a wide genomic interval and are generally not annotated. To overcome this challenge, we leveraged pan-cancer “enhancer-to-gene links” identified by the TCGA consortium based on the correlation of ATAC-Seq peaks to expression of nearby genes in matched tumors (24) (**Fig. 5A**). In every cancer type, the number of enhancer elements linked to each PRC2^+^-CGI gene was larger than that linked to each PRC2^−^-CGI gene (means of 2.9 v.s. 1.9, **Fig. 5B**). This suggested that enhancers play an important role in the regulation of PRC2^+^-CGI genes in cancer, as they do in normal development. We used HOMER to compare directly the frequency of TFBSs in PRC2^+^-v.s. PRC2^−^-CGI genes by setting one class as the foreground and the other as the background. Considering the distinct sequence contexts between promoter and enhancer regions (most notably, the high GC content and CpG density of CGI promoters), we performed separate analyses for promoters and enhancers. A notable pattern emerged from these reciprocal analyses: in promoter regions, PRC2^−^-CGI genes had enrichment for more TFBS motifs (80 v.s. 35 motifs for PRC2^+^-CGI genes, **Fig. 5C, left**). Conversely, in enhancer regions PRC2^+^-CGI class had enrichment for more motifs (64 v.s. 24 motifs, **Fig. 5C, right**).

**Fig. 5.**
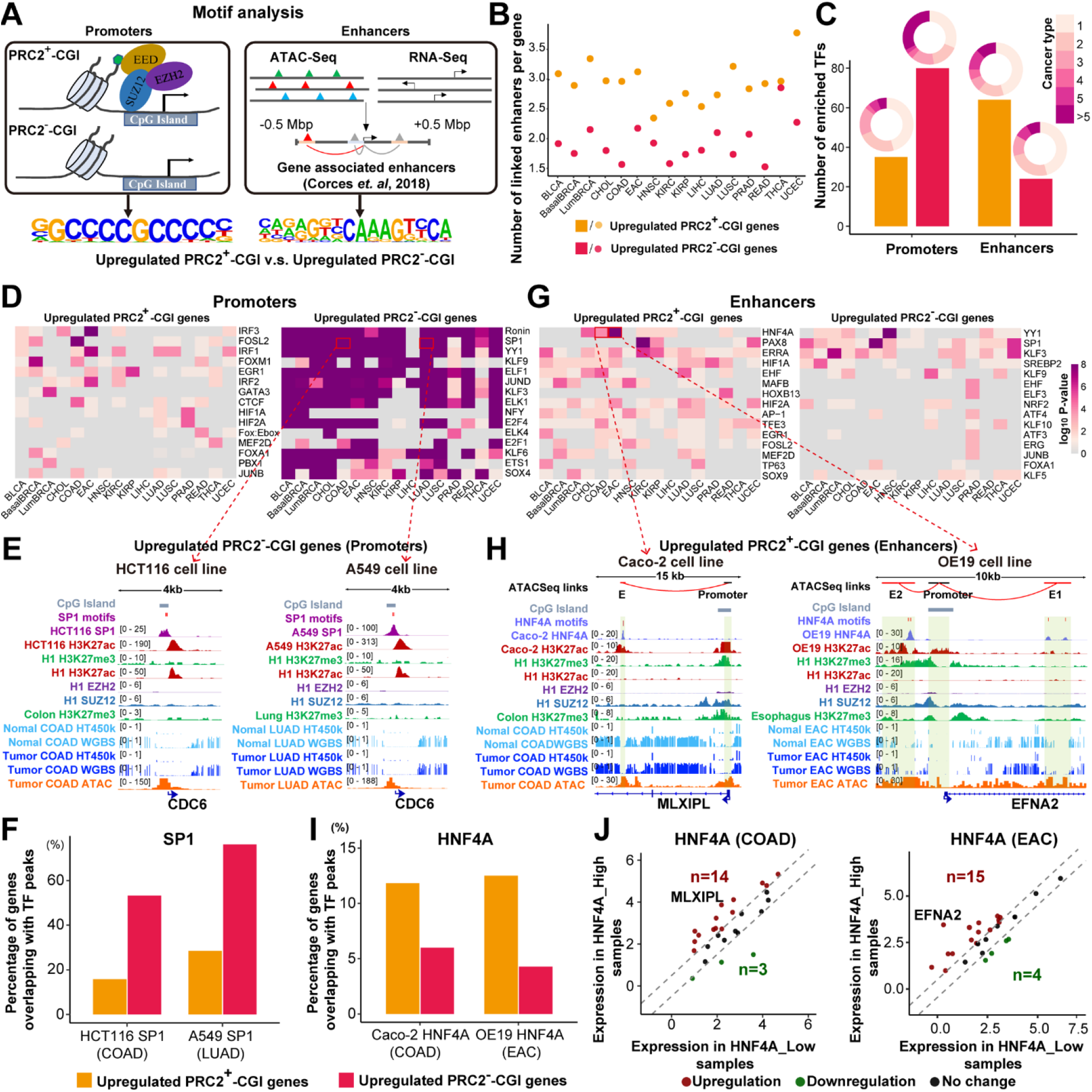
Upregulated PRC2^+^-CGI genes are linked to distal enhancers targeted by specific transcription factor binding sites (TFBSs). **(A)** Sequence motif enrichment analysis was performed for upregulated PRC2^+^-CGI and PRC2^−^-CGI genes using either promoter or enhancer regions. The linked enhancers are from “enhancer-to-gene links” defined by the TCGA consortium. **(B)** The number of linked enhancers per gene in both gene classes. **(C)** The number of significantly enriched TF motifs in both promoter and enhancer regions, with pie charts showing the number of cancer types found significant for each motif. **(D)** Top 15 enriched TFs identified in promoter regions. **(E)** IGV plots showing the promoter region of *CDC6* (a PRC2^−^-CGI gene) with predicted SP1 motifs and occupied by SP1 in HCT116 COAD cancer cells (left) and A549 LUAD cells (right) by ChIP-Seq from the ENCODE project. **(F)** TF ChIP-Seq of SP1 binding overlapping PRC2^+^-CGI vs. PRC2^−^-CGI promoters. **(G)** Top 15 enriched TFs identified in enhancer regions. **(H)** HNF4A-binding motif was predicted within distal enhancers for PRC2^+^-CGI genes *MLXIPL* in COAD and *EFNA2* in EAC, which was validated by HNF4A ChIP-Seq in COAD cells (Caco-2) and EAC cells (OE19). ChIP-Seq datasets were re-analyzed from GSE23436, GSE96069, E-MTAB-6858 and GSE132686. **(I)** TF ChIP-Seq of HNF4A binding overlapping PRC2^+^-CGI vs. PRC2^−^-CGI enhancers, from the same COAD and EAC dataset above. **(J)** Expression differences between TCGA HNF4A-high and HNF4A-low EAC/COAD tumors for the HNF4A target genes having enhancers overlapped by HNF4A in EAC or COAD cells (from panel **I**). High and low tumors were those in the upper and lower quintile of HNF4A expression.

We investigated the TF motifs that were most strongly enriched (top 15), starting with promoters (**Fig. 5D-E**). Promoter motifs were more abundant and ubiquitous in PRC2^−^-CGI class, while the few that were enriched in PRC2^+^-CGI genes tended to be more cancer-type specific. In fact, 32.5% of PRC2^−^-CGI promoter TFs were enriched in > 5 cancer types, compared to only 6% in the PRC2^+^-CGI group (colored rings in **Fig. 5C**). In PRC2^+^-CGI promoters, several known cancer-type-specific TFs were observed, such as FOXM1 in BasalBRCA (*36*) and GATA3 in LumBRCA (*37*); nevertheless, most TFs have not been reported in their corresponding cancer types. In contrast, in PRC2^−^-CGI promoters, a number of cell-cycle related TFs were significantly enriched across many cancer types, including SP1 (*38*), JUND (*39*), NFY (*40*), E2F4 (*41*), E2F1 (*41*) (**Fig. 5D**), supporting the results of our pathway enrichment analysis which also showed cell-cycle function among upregulated PRC2^−^-CGI genes. To validate these motif results, we compared published ChIP-Seq data to our predictions for the SP1 motif (the highest-ranking TF with available ChIP-Seq data). As predicted, in both HCT116 (COAD) and A549 (LUAD) cells, SP1-binding events were considerably more enriched in PRC2^−^- (53.2%-76.1%) than PRC2^+^-CGI promoters (15.8%-28.4%, **Fig. 5F**). As an example, the promoter of *CDC6* (a cell-cycle regulator) was directly occupied by SP1 in both cell lines (**Fig. 5E**). We also observed that multiple motifs of SP1 occurred within the same promoter region.

We next investigated the TF motifs that were most strongly enriched in enhancers (**Fig. 5G-H**). Enhancer motifs were more abundant among PRC2^+^-CGI genes, and these tended to have reported cancer-type-specific functions such as HNF4A in GI cancers (EAC, COAD, READ and CHOL) (*42*), TP63 in squamous cancers (LUSC, HNSC and a subset of BLCA) (*43*), PAX8 in KIRC (*44*) and UCEC (*45*), etc. In contrast, PRC2^−^-CGI genes had fewer motifs overall and fewer examples corresponding to established roles in cancer. We focused on two TFs enriched in PRC2^+^-CGI enhancers with publicly available ChIP-Seq data: HNF4A and TP63. HNF4A was most strongly enriched in GI cancers, and examples of HNF4A enhancer-binding are shown in these cell types (**Fig. 5H;** no CHOL or READ cell lines had available HNF4A ChIP-Seq data). TP63 was most strongly enriched in LUSC, and an example of TP63 enhancer-binding in LUSC is shown for the *TP73* gene (**Fig. S8A**). We further validated that HNF4A enhancer-binding specifically targeted PRC2^+^-CGI genes by calculating the number of genes with linked enhancers covered by HNF4A ChIP-Seq peaks (11.8%-12.5%), and comparing it to the number of PRC2^−^- CGI genes (4.3%-6.0%) (**Fig. 5I**). A very similar trend was observed for TP63 (**Fig. S8B**).

We next explored the correlation between the expression of HNF4A and that of its PRC2^+^-CGI target genes (n=27 in COAD, n=28 in EAC; **Fig. 5I**) across TCGA COAD and EAC tumors. We binned tumors into HNF4A-high and HNF4A-low groups based on the top and bottom quintiles of HNF4A expression, and plotted the levels of the target genes (**Fig. 5J**). More than half of these PRC2^+^-CGI genes (14/27 for COAD; 15/28 for EAC) had higher expression in the HNF4A-high samples, and only 3-4 genes were lower. A highly similar trend of TP63 targets was observed in LUSC tumors (**Fig. S8C**).

### HNF4A regulates PRC2^+^-CGI target genes through activation of distal enhancers

In order to better illuminate functional mechanisms, we continued to focus on the 28 PRC2^+^- CGI genes that were upregulated in EAC and had ChIP-Seq HNF4A-binding sites in linked enhancers. We first re-analyzed public ATAC-Seq chromatin accessibility datasets of nonmalignant esophageal epithelium and EAC tumors, focusing on these 28 PRC2^+^-CGI genes and their 33 linked enhancers. In nonmalignant esophageal epithelium, only 4 out of the 33 linked enhancers had ATAC-Seq peaks (**Fig. 6A, right**). In EAC tumors, 21 of the remaining 29 enhancers gained peaks (**Fig. 6A, right**). The independent TCGA ATAC-Seq dataset (**Fig. 6B**) did not contain nonmalignant samples, but had EAC tumors which we could compare to the other molecular subtype, ESCC. In our analysis, 23 of the 33 HNF4A-occupied enhancers had significantly higher ATAC-Seq signals in EAC, and none had lower (**Fig. 6B**). We next re-analyzed ATAC-Seq following ectopic expression of HNF4A in normal esophageal cells (**Fig. 6C**). In agreement with patient samples, 31 of the 33 HNF4A-binding enhancers were inaccessible in normal esophageal cells (Het1A), and about half (16/31) became accessible upon HNF4A overexpression.

**Fig. 6.**
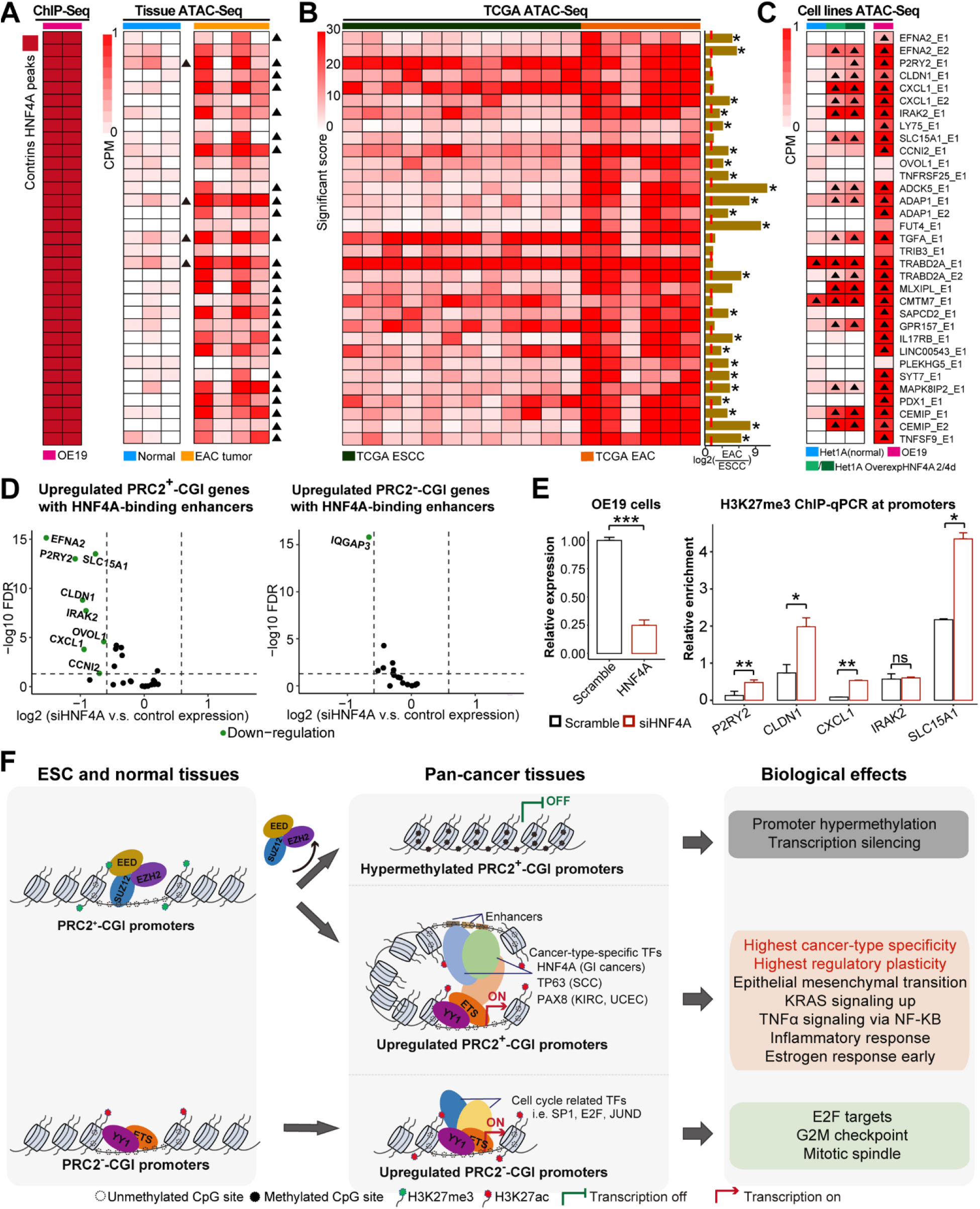
Experimental validation of HNF4A as an upstream regulator of PRC2^+^-CGI genes through activating distal enhancers. **(A-C)** Focusing on the 33 HNF4A-binding enhancers, **(A)** shows HNF4A ChIP-Seq peaks (left) and ATAC-Seq peaks in EAC normal and tumor tissues (right), and **(B)** TCGA ESCC and EAC tumor tissues as well as **(C)** EAC cell lines. The ATAC-Seq signals in **(A)** are normalized with CPM method and those called as peaks are marked with a triangle. TCGA ATAC-Seq signals are normalized by the TCGA consortium and the fold change of ATAC-Seq signals between EAC and ESCC samples are calculated. Those genes with significant increase in EAC tumors were marked with a star (student t-test, p value<0.05). **(D)** In EAC cells (OE19) with HNF4A knockdown, volcano plots showing expression changes of either PRC2^+^-CGI (left) or PRC2^−^-CGI genes (right) having HNF4A-binding enhancers. **(E)** Promoter H3K27me3 signals were measured by ChIP-qPCR in both scramble and siHNF4A OE19 cells. **(F)** A summary graph illustrating the cancer-specific deregulation of both PRC2^+^-CGI and PRC2^−^-CGI genes, the underlying molecular mechanisms and the biological implications.

The datasets described above demonstrate the direct regulation of these target genes by the interaction of HNF4A with the linked enhancers in an EAC-specific manner. To further validate this finding, we re-analyzed the HNF4A-wildtype and HNF4A-knockdown RNA-Seq datasets in an EAC cell line (OE19). We found that 28.5% (8/28) of the HNF4A-linked PRC2^+^-CGI genes were down-regulated (**Fig. 6D, left**), compared to the background level for all PRC2^+^-CGI genes of 3.5% (152/4,378). 6 of the 8 downregulated genes also gained ATAC-Seq peaks in the HNF4A overexpression assay in Het1A cells (**Fig. 6C, left**). Additionally, this regulation appeared to be PRC2^+^-CGI specific - only 1 out of 17 PRC2^−^-CGI genes (5.9%) overlapping with HNF4A ChIP-Seq was downregulated in the knockdown (**Fig. 6D, right**).

Because H3K27me3 data was unavailable for any of these cell types other than normal esophagus, we performed promoter H3K27me3 ChIP-qPCR in OE19 HNF4A-wildtype and HNF4A-knockdown cells. We performed this assay for all 5 genes that were downregulated upon HNF4A-knockdown and gained enhancer accessibility under HNF4A-overexpression (the 6^th^ such gene, *EFNA2*, was not determined because no optimal primer could be designed for its promoter region). Four of these five gene promoters showed gain of H3K27me3 signal in the knockdown of HNF4A (**Fig. 6E**). Taken together, these results characterize HNF4A as an upstream regulator of PRC2^+^-CGI genes in EAC by activating distal enhancers and removing PRC2-associated H3K27me3 at promoters.

## Discussion

Most PRC2-occupied promoters overlap CpG Islands and are known to be prone to *de novo* DNA hypermethylation and transcriptional repression in cancer, but few studies have looked systematically at expression changes in these PRC2^+^-CGI promoters and the larger class of PRC2^−^-CGI promoters (*18–20*). Here we comprehensively investigated cancer-associated deregulation of all CGI promoter genes across pan-cancer samples, revealing regulatory similarities and differences between these two classes of genes. Consistent with prior findings (*1, 4*), we showed that many PRC2^+^-CGI genes were commonly hypermethylated and downregulated in most cancers, affecting 2,274/4,378 genes across 16 cancer types. Unexpectedly, we also found a large class of PRC2^+^-CGI genes (1,543/4,378) to be upregulated in one or more cancer types. Among these upregulated PRC2^+^-CGI genes, we found many well-defined oncogenes and tumor-promoting factors such as MYB, TWIST1, SYK, TEAD4, FOXC1, and FGFR3 (**Table S3B**). Previous studies in normal cells have demonstrated that PRC2^+^-CGI promoters are unmethylated (*11*), with limited chromatin accessibility (*46*) and weak transcriptional activity (*17*). In tumors, our analysis showed that unlike hypermethylated PRC2^+^-CGI promoters, upregulated PRC2^+^-CGI promoters gain accessibility and the H3K27ac mark (illustrated in **Fig. 6F**). While upregulated PRC2^−^-CGI promoters also gained these active characteristics, they were much more likely to start with high baseline levels of these features in nonmalignant tissues. This helps explain why upregulated PRC2^−^-CGI genes had higher absolute expression in tumors, but PRC2^+^-CGI genes had higher fold-change differences from normal tissues (median of 3.5 for PRC2^+^-CGI v.s. 2.9 for PRC2^−^-CGI genes).

Among our most intriguing findings was the high degree of cancer-type specificity in expression of upregulated PRC2^+^-CGI genes, which was markedly higher than either upregulated PRC2^−^- CGI or hypermethylated PRC2^+^-CGI class. This property allowed for better clustering of cancer types and subtypes using the PRC2^+^-CGI class than either of the other classes. Interestingly, nearly half (762/1,543) of upregulated PRC2^+^-CGI genes were also observed to be hypermethylated in other cancer types, including some known tumor suppressors, including *DKK1*, *NFGR*, *PRICKLE1*. For example, *DKK1* was hypermethylated in LumBRCA, whereas it became upregulated in both BasalBRCA and LUSC. *DLX5* was similarly hypermethylated in LumBRCA, but it was upregulated in multiple squamous type cancers (**Table S3**) as detailed in a new functional study of this gene (Manuscript under revision in this Journal). These findings suggest a bifurcated chromatin re-configuration of many PRC2^+^-CGI genes (“plastic” genes) during tumorigenesis, dependent on different transcriptional programs and TF activities in different cell types. This regulatory plasticity is not entirely surprising given the disproportional role PRC2^+^-CGI genes play in normal patterning and development. The regulatory complexity of PRC2^+^-CGI genes in cancer was also evident from the types of biological pathways we identified among these genes compared to other CGI genes, including important cancer-related pathways such as EMT, KRAS, and TNFα signaling and inflammatory response. Indeed, these pathways were often cancer-type specific.

Functionally, upregulated PRC2^+^- and PRC2^−^-CGI genes control distinct sets of Hallmark pathways in cancer (**Fig. 4A-B**). Specifically, upregulated PRC2^−^-CGI genes are highly enriched in cell-cycle pathways invariably across different cancer types. This is also consistent with the motif enrichment result that enriched TFs in PRC2^−^-CGI class are strongly overlapped in different cancer types and are associated with cell-cycle pathways, such as SP1, JUND, NFY, E2F4 and E2F1. Furthermore, over half of PRC2^−^-CGI genes in cell-cycle pathways are shared across cancer types, highlighting the common activation of cell-cycle-related PRC2^−^-CGI genes in cancer. In comparison, a completely different set of biological pathways are enriched in upregulated PRC2^+^-CGI genes, such as KRAS, EMT, and TNFα signaling and inflammatory response pathways. We performed genetic manipulations and confirmed the role of PRC2^+^-CGI genes in the EMT pathway. In addition, our finding of the involvement of a subset of PRC2^+^-CGI genes in immunologically “hot” tumors may have important implications for immune-checkpoint blockade therapy, especially given that the enzymatic activity of PRC2 can be pharmacologically targeted (e.g., by EZH2 inhibitors). Indeed, consistent with this notion, recent studies have shown that EZH2 inhibition leads to hightented anti-cancer immunity and synergizes with immune-checkpoint blockade therapy in different cancer types (*47, 48*).

Our conclusions above regarding cancer-type-specificity strongly suggest that the upregulated PRC2^+^-CGI genes are controlled by distal enhancers, which govern cell-type-specific expression programs and have been shown to regulate the PRC2 status of linked promoters (*49*). By using cancer type-specific enhancer links from the TCGA ATAC-Seq project (*24*) combined with TFBS motif analysis, we were able to show that PRC2^+^-CGI genes were predominantly linked to specific TFBSs in distal enhancers, whereas PRC2^−^-CGI genes were linked to TFBSs in promoters (as illustrated in **Fig. 6F**). Unsurprisingly, the TFs whose binding sites were enriched in PRC2^−^-CGI promoters exhibited ubiquitous expression pattern across cancer types, whereas the enhancer-linked TFBSs were enriched in specific cancer types. We functionally validated a few of these TF/target-gene relationships using publicly available ATAC-Seq and ChIP-Seq datasets as well as genetic perturbations. For HNF4A, a master regulator of GI cancers (*42*), loss of function in cancer cells led to gain of the H3K27me3 mark at promoters and reduced expression of genes linked to HNF4A-occupied enhancers. This mode of action in cancer is consistent with the model proposed in Taberlay *et al.* for PRC2^+^-CGI genes during normal development (*49*). While this mode of activation appears to be prevalent in cancer based on our analysis, additional layers of deregulation of these genes may be caused by genetic disruption of PRC2 proteins themselves, given the discovery of both loss-of-function and gain-of-function mutations of PRC2 complex (particularly *EZH2*) in cancer (*50*).

In summary, we have systematically investigated the cancer-specific deregulation of different classes of CGI promoters, which together make up ~70% of all human promoters. We identified bifurcated deregulation of PRC2^+^-CGI genes, leading to either hypermethylation-associated gene silencing or transcriptional activation depending on the cancer type. The PRC2^+^-CGI genes that become silenced have been well-studied, but those that become activated have not, and appear to play an important role in pathways such as EMT and TNFα associated inflammatory response in cancer. Finally, we show that many of these activating events are controlled by the activity of specific TFs in distal enhancers linked to these genes. These data together advance our mechanistic understanding of the chromatin regulation of these different gene categories in cancer, while providing a comprehensive catalog of candidate cancer-associated genes for future investigation.

## Materials and Methods

### Data sources

The TCGAbiolinks package (*51*) was used to download the sample information, mRNA expression (RNA-Seq level 3 data) and DNA methylation (Illumina HumanMethylation450 array) data of 33 types of cancers (n=10,528) from the TCGA project. All the TCGA data were downloaded in GDC v16.0. Considering the distinct biology between established cancer subtypes (including esophageal adenocarcinoma v.s. squamous cancer, breast luminal v.s. basal cancer, lung adenocarcinoma v.s. squamous cancer), they were analyzed as distinct disease subtypes. To ensure the statistical power for comparing nonmalignant and tumor tissues, cancer types with fewer than 5 nonmalignant samples were excluded, resulting in 16 cancer types available for analyses (**Table S1**). For statistical tests, each tumor type was analyzed independently to avoid potential batch effects between TCGA disease projects. ATAC-Seq (Assay for Transposase-Accessible Chromatin using Sequencing) data of TCGA samples and pan-cancer “enhancer-to-gene” links were obtained from a recent TCGA publication (*24*).

The following additional datasets were collected: H3K27ac ChIP-Seq in nonmalignant colonic crypts and primary colon cancer cells (GSE77737), H3K27ac ChIP-Seq in nonmalignant and tumor samples of kidney renal clear cell carcinoma (KIRC) from GSE86095, HNF4A ChIP-Seq in OE19 (E-MTAB-6858) and Caco-2 (GSE23436) cell lines, TP63 ChIP-Seq in HCC95 cell line (GSE46837), SP1 and JUND ChIP-Seq in HCT116 and A549 cell lines (ENCODE), H3K27ac ChIP-Seq in OE19 (GSE132686), HCC95 (GSE66992), HCT116 (ENCODE), Caco-2 (GSE96069) and A549 (ENCODE) cell lines. RNA-Seq of HNF4A knockdown, ATAC-Seq of nonmalignant esophageal epithelium, EAC tissues, normal esophageal cells (HET1A) and OE19 tumor cells were downloaded from E-MTAB-6756, E-MTAB-6751 and E-MTAB-6931. RNA-Seq datasets from pre-treatment tumors with anti-PD-1 monotherapy in KIRC were obtained from Miao *et. al.* (*35*). We also retrieved the mRNA expression data of basal breast cancer cell lines from the Cancer Cell Line Encyclopedia (CCLE). Annotation of CGI regions was downloaded from UCSC website (http://hgdownload.soe.ucsc.edu/goldenPath/hg38/database/).

### Curation of CGI promoters

We extracted all transcription start sites (TSSs) from the GENCODE basic annotation file (version 31). Because Illumina HumanMethylation450 array (which was used by TCGA) contains FANTOM4-annotated promoters (*52*), TSSs from FANTOM4 were also integrated in our study. All the genome coordinates were converted to hg38 using the UCSC LiftOver function (https://genome.ucsc.edu/cgi-bin/hgLiftOver). We extracted the promoter regions from 250bp upstream to 500bp downstream (−250bp ~ +500bp) of the TSSs. Promoters which are not covered by any methylation probes were excluded for further analyses. The average β values were calculated to represent the methylation level of each promoter. We merged neighboring promoters covered by the same methylation probes and excluded those on either Y chromosome or mitochondria. Next, we used the GENCODE comprehensive annotation file (version 31) for the annotation of FANTOM4 promoters via bedtools intersect function (https://bedtools.readthedocs.io/en/latest/). Finally, only promoters overlapping with CGI regions (that is, CGI promoters) were retained for further analyses, based on the CpG Island track from the UCSC browser (Gardener-Garden criteria) (**Fig. S1**).

### Identification of PRC2^+^-CGI genes in ESC (H1 cells) and normal tissues

H3K27me3 and H3K27ac ChIP-Seq profiles in both ESCs (H1) and normal tissues (colonic mucosa, lung, breast epithelium, rectum, esophagus, uterus and liver) were obtained from the combined NIH RoadMap / ENCODE data repository. Based on the 15-state epigenomic model established by Ernst *et. al.* in ESCs (*21*), we first obtained H3K27me3-positive regions by retrieving “State 10: Bivalent/Poised TSS”, “State 11: Flanking Bivalent TSS/Enhancers” and “State 13: Repressed PolyComb”. From these H3K27me3-positive regions, we identified PRC2-occupied regions by requiring them to have either EZH2- or SUZ12-binding (ChIP-Seq datasets were downloaded from ENCODE project).

Because neither EZH2- nor SUZ12-binding was available in normal tissues, we next analyzed both the expression and epigenomic states of ESC PRC2^+^-CGI genes in normal tissues. An FPKM value of 4 in TCGA normal tissues readily separated PRC2^+^-CGI genes with divergent H3K27ac levels: PRC2^+^-CGI genes with FPKM < 4 had considerably lower H3K27ac signals than those with FPKM >= 4 (**Fig. S2A**). Furthermore, we confirmed that PRC2^+^-CGI genes with FPKM < 4 had much higher H3K27me3 level than those with FPKM >= 4 (**Fig. S2B**). These results demonstrate that a major subset of PRC2^+^-CGI genes in ESCs (FPKM < 4) had conserved PRC2-occupancy in normal tissues, and this subset of PRC2^+^-CGI genes were selected for further analysis (**Fig. 1A**).

### Classification of upregulated PRC2^+^-CGI and PRC2^−^-CGI genes in cancer

Based on the raw read count matrices downloaded from TCGA, all expressed genes with i) read counts > 0 in more than 80% both nonmalignant and tumor tissues of each cancer type, and ii) FPKM value > 1 in more than half of tumor samples, were used for differential expression analysis (**Fig. 1A**). DESeq2 package (*53*) was applied and those with adjusted P value < 0.05 and log2 fold change (tumor v.s. nonmalignant) > 1 were considered as upregulated genes in cancer for both PRC2^+^- and PRC2^−^-CGI classes.

### Classification of hypermethylated PRC2^+^-CGI genes in cancer

To identify PRC2^+^-CGI genes with DNA hypermethylated promoters in cancer, we applied criteria based on those developed by the TCGA consortium (*23*): methylation β values below 0.2 in more than 90% nonmalignant tissues and above 0.3 in over 15% tumor samples (**Fig. 1A**). The resulting genes were additionally required to have no significant upregulation in tumor than nonmalignant samples (**Fig. 1A**).

### Cell culture

Breast cancer cell lines HS578T and MDAMB436 were cultured in Dulbecco’s modified Eagle medium (DMEM) (Thermo Fisher Scientific, USA) and EAC cell line OE19 was cultured in RPMI-1640 medium (Thermo Fisher Scientific, USA). All the medium was supplemented with 10% fetal bovine serum (FBS) (Omega Scientific, USA) and 100 U/ml penicillin and 100 mg/ml streptomycin (Thermo Fisher Scientific, USA).

**Additional methods are described in the Supplementary Materials.**

## Supplementary Materials

**Supplementary Methods and Figures.docx**

**Table S1. TCGA nonmalignant and tumor samples used in the present study**

**Table S2. Numbers of gene and promoter of different CGI categories across cancer types**

**Table S3. Different groups of CGI genes across different cancer types. (A)** Hypmethylated PRC2^+^-CGI genes, **(B)** Upregulated PRC2^+^-CGI genes and **(C)** Upregulated PRC2^−^-CGI genes.

**Table S4. siRNA and primers used in this study**

## Author contributions

D.-C.L. and B.P.B conceived and devised the study. D.-C.L., B.P.B., Y.Y.Z and G.W.H. designed experiments and analyses. Y.Y.Z, T.C.S and Q.Y. performed bioinformatics and statistical analysis. G.W.H. performed the experiments. Y.Y.Z., B.P.B. and D.-C.L. analyzed the data. H.P.K, B.P.B, D.-C.L. supervised the research. D.-C.L. wrote the manuscript with assistance from Y.Z., H.P.K., and B.P.B.

