## Supplementary Methods and Figures for "A pan-cancer analysis of CpG Island gene regulation reveals extensive plasticity within Polycomb targets"

#### Data sources

The following datasets were collected: H3K27ac ChIP-Seq in nonmalignant colonic crypts and primary colon cancer cells (GSE7773, 7 tumor samples were removed as the number of reads is smaller than 10M) (*1*), H3K27ac ChIP-Seq in nonmalignant and tumor samples of kidney renal clear cell carcinoma (KIRC) from GSE86095 (3 pairs of nonmaligant and tumor samples were removed due to insufficient number of peaks) (*2*), HNF4A ChIP-Seq in OE19 (E-MTAB-6858) (*3*) and Caco-2 (GSE23436) (*4*) cell lines, TP63 ChIP-Seq in HCC95 cell line (GSE46837) (*5*), SP1 and JUND ChIP-Seq in HCT116 and A549 cell lines (ENCODE), H3K27ac ChIP-Seq in OE19 (GSE132686), HCC95 (GSE66992) (*6*), HCT116 (ENCODE), Caco-2 (GSE96069) (*7*) and A549 (ENCODE) cell lines. RNA-Seq of HNF4A knockdown, ATAC-Seq of nonmalignant esophageal epithelium，EAC tissues, normal esophageal cells (HET1A) and OE19 tumor cell line were downloaded from E-MTAB-6756, E-MTAB-6751 and E-MTAB-6931 (*8*). RNA-Seq datasets from pre-treatment tumors with anti-PD-1 monotherapy in KIRC were obtained from Miao *et. al*. (*9*) We also retrieved the mRNA expression data of basal breast cancer cell lines from the Cancer Cell Line Encyclopedia (CCLE) (*10*). Annotation of CGI regions was downloaded from UCSC website (http://hgdownload.soe.ucsc.edu/goldenPath/hg38/database/).

#### ChIP-Seq data analysis

Raw reads with length shorter than 51bp were aligned to GRCh38 (ENSEMBL release 84) using Bowtie with "--best --chunkmbs 200" options (*11*). Bowtie2 was applied for those reads longer than 51bp with the "--sensitive" parameters (*12*). Then the uniquely mapped reads were extracted and sorted by SAMtools program using "-f 2 -q 10" options (*13*). PCR duplicates and blacklist regions were removed by Picard MarkDuplicates tool (http://broadinstitute.github.io/picard/) and bedtools, respectively. ChIP-Seq peaks were called using MACS2 (Model-Based Analysis of ChIP-Seq, version 2.1.2) (*14*) with the default parameters for TFs and “-q 0.01 --extsize=146 --nomodel -B” options for H3K27ac. Reads were extended at default setting and normalized at -log10 of the Poisson p-value by MACS2 bdgcmp command using “-m ppois” options. BedGraphToBigWig tool was used to generate the BigWig files (*15*).

**ATAC-Seq data analysis**

ATAC-Seq data were analyzed using the published pipeline (*16*). Bowtie2 was applied for pre-alignments to filter out reads that align to repetitive regions using “-k 1 -D 20 -R 3 -N 1 -L 20 -i S,1,0.50 -X 2000 –rg-id” parameters. For the remaining reads, Bowtie2 was used to map to GRCh38 with “--very-sensitive -X 2000 --rg-id” options. Then, the SAMtools program was applied to sort and extract uniquely mapped reads, followed by the removal of PCR duplicates. Next, ATAC-Seq peaks were identified using MACS2 with “--shift -75 --extsize 150 -B --nomodel --call-summits --keep-dup all -q 0.01” parameters.

**RNA-Seq analysis**

For RNA-Seq datasets of control and siHNF4A in OE19 cell line, 75bp paired-end reads were mapped to GRCh38 using HISAT2 (version 2.0.4) (*17*) and counted by htseq-count program (version 0.11.2) with default parameters. Differentially expressed genes were identified by the DESeq2 package with adjusted P value < 0.05 and absolute log2 fold change (siHNF4A v.s. control) > 0.5.

**Principal component analysis (PCA)**

PCA was performed using the R prcomp function and point plots were generated by the ggplot2 package.

**Hallmark pathway enrichment analysis**

Cancer hallmark gene sets were obtained from MSigDB (*18*). Upregulated $PRC2^{+}$- and $PRC2^{-}$-CGI genes were used as the foreground and background, respectively. Then the population test was performed on these two groups and the enriched hallmarks with p value <0.01 were identified. To reveal the functional difference between these two groups of genes, we switched the foreground with background and repeated the same test.

##### TF-binding sequence motif enrichment analysis

For the two groups of upregulated CGI promoters, sequence motif analyses were first performed using their promoter regions to identify potential TF-binding sequences through HOMER findMotifsGenome.pl script (*19*) using either one group as the foreground and the other as the background. To study enhancer regions, we used the pan-cancer "enhancer-to-gene" links and performed the same motif analyses. Considering that TFs from the same TF family can recognize identical binding sequences (such as GATA and SOX families), we reserved those enriched TFs with FPKM > 10 in the corresponding cancer types.

##### siRNA transfection and real-time RT-PCR

siRNA oligos were synthesized by IGEbio (IGEBio, China) and transfected into cells by Lipofectamine RNAiMAX (Thermo Fisher Scientific, USA). The cells were collected and RNA was extracted 72h post-transfection. Real-time PCR was performed by using Power SYBR Green Master Mix (Thermo Fisher Scientific, USA). All the sequences for siRNAs and the primers for PCR were shown in **Supplementary Table S4**.

##### Chromatin immunoprecipitation (ChIP)

Chromatin immunoprecipitation was performed as previously described (*20*) with slight modification. Briefly, OE19 cells were cross-linked with formaldehyde at final concentration of 1.42% for 15 min at room temperature and followed by 125 mM glycine for 5 min. The cells were collected and lysed with lysis buffer [150 mM NaCl, 50 mM Tris-HCl (pH 7.5), 5 mM EDTA, NP-40 (0.5% vol/vol), Triton X-100 (1.0% vol/vol)] and the nuclear pellet was collected after centrifuge by 10,000 g, 1min at 4 ℃ and resuspended in 1ml shearing buffer [(20% SDS, 0.5 M EDTA (pH 8.0), 1 M Tris (pH 8.0)] for sonication. Cell debris was removed by centrifuge and the supernatant was diluted with 5-fold volume dilution buffer [(20% SDS, 0.5 M EDTA (pH 8.0), 1 M Tris (pH 8.0), 1% Triton X-100, 5M NaCl]. 2% of the lysate was used as input and the rest was incubated with an anti-H3K27Me3 antibody (Abcam Biotechnology, UK) at 4 ℃ overnight. The next day, protein G-coupled magnetic beads (Thermo Fisher Scientific, USA) were added to pulldown protein-DNA complex. After washing the precipitated complex with lysis buffer, the complex was eluted with buffer (150 μl 0.5 M NaHCO3, 50 μl 20% SDS) and subsequently subject to de-crosslink with supplement of 8 μl 5 M NaCl at 65 ℃ overnight. DNA was extracted after the complex was treated with RNase A and proteinase K (Thermo Fisher Scientific, USA) and used for real-time PCR. The primers for PCR are provided in **Supplementary Table S4**.

#### Supplementary Methods Reference


1. A. J. Cohen, A. Saiakhova, O. Corradin, J. M. Luppino, K. Lovrenert, C. F. Bartels, J. J. Morrow, S. C. Mack, G. Dhillon, L. Beard, L. Myeroff, M. F. Kalady, J. Willis, J. E. Bradner, R. A. Keri, N. A. Berger, S. M. Pruett-Miller, S. D. Markowitz, P. C. Scacheri, Hotspots of aberrant enhancer activity punctuate the colorectal cancer epigenome. *Nat. Commun.* **8**, 14400 (2017).
2. X. Yao, J. Tan, K. J. Lim, J. Koh, W. F. Ooi, Z. Li, D. Huang, M. Xing, Y. S. Chan, J. Z. Qu, S. T. Tay, G. Wijaya, Y. N. Lam, J. H. Hong, A. P. Lee-Lim, P. Guan, M. S. W. Ng, C. Z. He, J. S. Lin, T. Nandi, A. Qamra, C. Xu, S. S. Myint, J. O. J. Davies, J. Y. Goh, G. Loh, B. C. Tan, S. G. Rozen, Q. Yu, I. B. H. Tan, C. W. S. Cheng, S. Li, K. T. E. Chang, P. H. Tan, D. L. Silver, A. Lezhava, G. Steger, J. R. Hughes, B. T. Teh, P. Tan, Deficiency Drives Enhancer Activation of Oncogenes in Clear Cell Renal Cell Carcinoma. *Cancer Discov.* **7**, 1284–1305 (2017).
3. C. Rogerson, E. Britton, S. Withey, N. Hanley, Y. S. Ang, A. D. Sharrocks, Identification of a primitive intestinal transcription factor network shared between esophageal adenocarcinoma and its precancerous precursor state. *Genome Res.* **29**, 723–736 (2019).
4. M. P. Verzi, H. Shin, H. H. He, R. Sulahian, C. A. Meyer, R. K. Montgomery, J. C. Fleet, M. Brown, X. S. Liu, R. A. Shivdasani, Differentiation-specific histone modifications reveal dynamic chromatin interactions and partners for the intestinal transcription factor CDX2. *Dev. Cell*. **19**, 713–726 (2010).
5. H. Watanabe, Q. Ma, S. Peng, G. Adelmant, D. Swain, W. Song, C. Fox, J. M. Francis, C. S. Pedamallu, D. S. DeLuca, A. N. Brooks, S. Wang, J. Que, A. K.

Rustgi, K.-K. Wong, K. L. Ligon, X. S. Liu, J. A. Marto, M. Meyerson, A. J. Bass, SOX2 and p63 colocalize at genetic loci in squamous cell carcinomas. *J. Clin. Invest.* **124**, 1636–1645 (2014).

6. X. Zhang, P. S. Choi, J. M. Francis, M. Imielinski, H. Watanabe, A. D. Cherniack, M. Meyerson, Identification of focally amplified lineage-specific super-enhancers in human epithelial cancers. *Nat. Genet.* **48**, 176–182 (2016).

7. Y. Nakamura, N. Hattori, N. Iida, S. Yamashita, A. Mori, K. Kimura, T. Yoshino, T. Ushijima, Targeting of super-enhancers and mutant BRAF can suppress growth of BRAF-mutant colon cancer cells via repression of MAPK signaling pathway. *Cancer Lett.* **402**, 100–109 (2017).

8. C. Rogerson, E. Britton, S. Withey, N. Hanley, Y. S. Ang, A. D. Sharrocks, Identification of a primitive intestinal transcription factor network shared between esophageal adenocarcinoma and its precancerous precursor state. *Genome Res.* **29**, 723–736 (2019).

9. D. Miao, C. A. Margolis, W. Gao, M. H. Voss, W. Li, D. J. Martini, C. Norton, D. Bossé, S. M. Wankowicz, D. Cullen, C. Horak, M. Wind-Rotolo, A. Tracy, M. Giannakis, F. S. Hodi, C. G. Drake, M. W. Ball, M. E. Allaf, A. Snyder, M. D. Hellmann, T. Ho, R. J. Motzer, S. Signoretti, W. G. Kaelin Jr, T. K. Choueiri, E. M. Van Allen, Genomic correlates of response to immune checkpoint therapies in clear cell renal cell carcinoma. *Science*. **359**, 801–806 (2018).

10. M. Ghandi, F. W. Huang, J. Jané-Valbuena, G. V. Kryukov, C. C. Lo, E. R. McDonald 3rd, J. Barretina, E. T. Gelfand, C. M. Bielski, H. Li, K. Hu, A. Y. Andreev-Drakhlin, J. Kim, J. M. Hess, B. J. Haas, F. Aguet, B. A. Weir, M. V. Rothberg, B. R. Paolella, M. S. Lawrence, R. Akbani, Y. Lu, H. L. Tiv, P. C. Gokhale, A. de Weck, A. A. Mansour, C. Oh, J. Shih, K. Hadi, Y. Rosen, J. Bistline, K. Venkatesan, A. Reddy, D. Sonkin, M. Liu, J. Lehar, J. M. Korn, D. A. Porter, M. D. Jones, J. Golji, G. Caponigro, J. E. Taylor, C. M. Dunning, A. L. Creech, A. C. Warren, J. M. McFarland, M. Zamanighomi, A. Kauffmann, N. Stransky, M. Imielinski, Y. E. Maruvka, A. D. Cherniack, A. Tsherniak, F. Vazquez, J. D. Jaffe, A. A. Lane, D. M. Weinstock, C. M. Johannessen, M. P. Morrissey, F. Stegmeier, R. Schlegel, W. C. Hahn, G. Getz, G. B. Mills, J. S. Boehm, T. R. Golub, L. A. Garraway, W. R. Sellers, Next-generation characterization of the Cancer Cell Line Encyclopedia. *Nature*. **569**, 503–508 (2019).

11. B. Langmead, C. Trapnell, M. Pop, S. L. Salzberg, Ultrafast and memory-efficient alignment of short DNA sequences to the human genome. *Genome Biol.* **10**, R25 (2009).

12. B. Langmead, S. L. Salzberg, Fast gapped-read alignment with Bowtie 2. *Nat. Methods*. **9**, 357–359 (2012).

13. H. Li, B. Handsaker, A. Wysoker, T. Fennell, J. Ruan, N. Homer, G. Marth, G. Abecasis, R. Durbin, 1000 Genome Project Data Processing Subgroup, The

Sequence Alignment/Map format and SAMtools. *Bioinformatics*. **25**, 2078–2079 (2009).

### Supplementary Figures

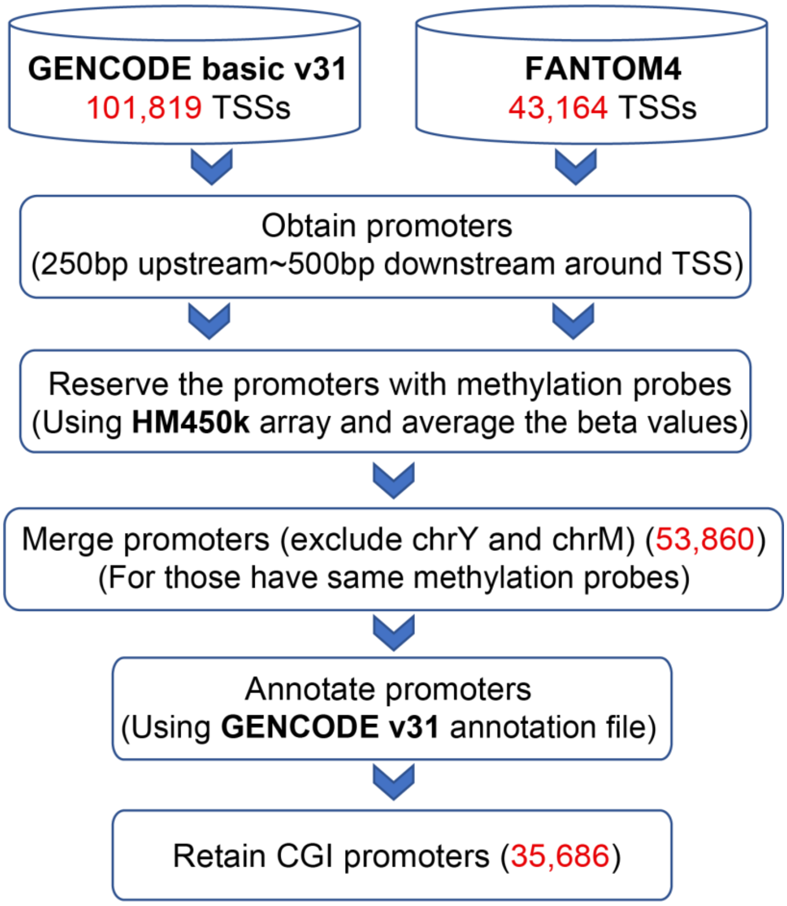


**Fig. S1. The workflow of obtaining the CGI promoters**

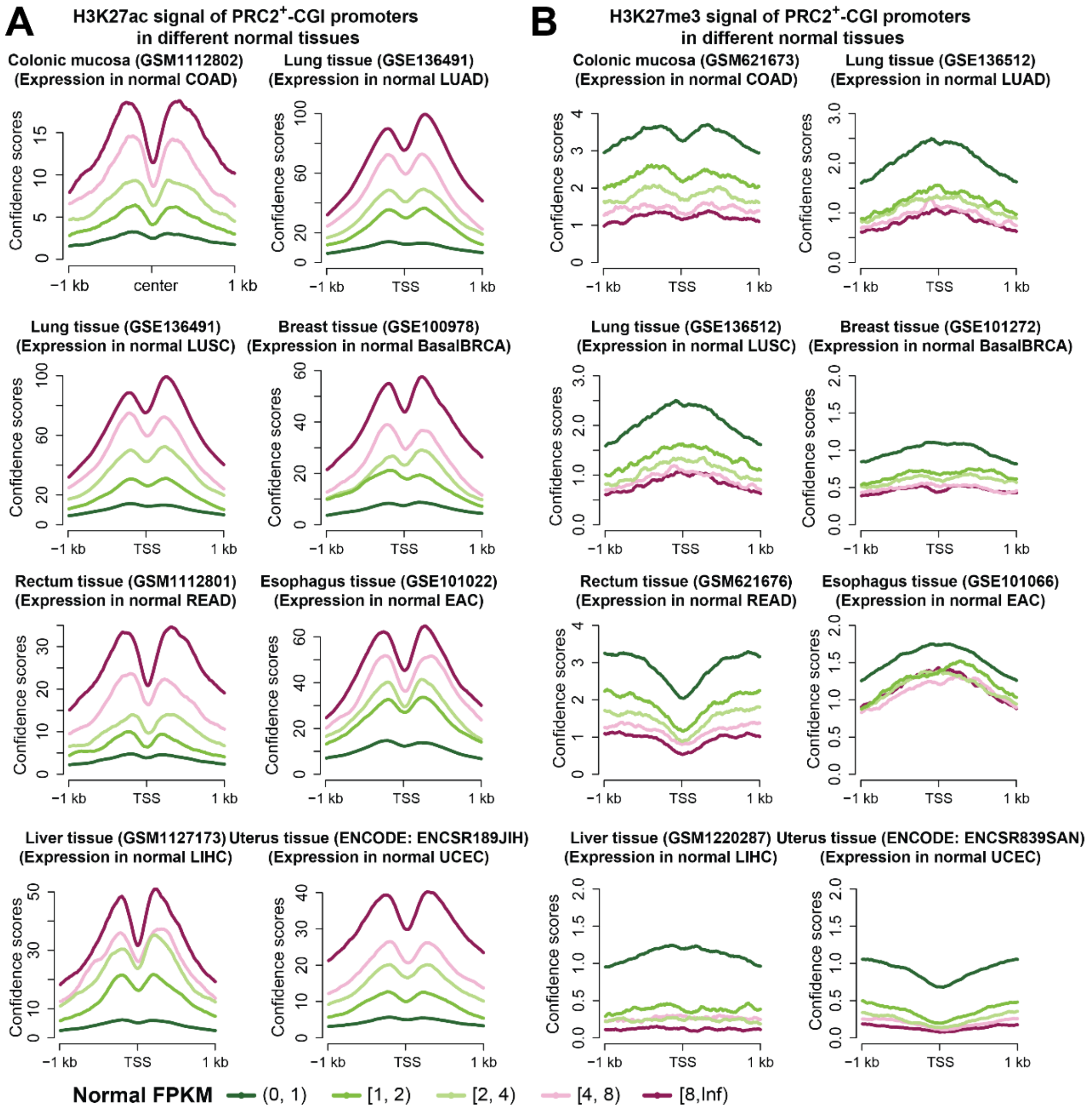


**Fig. S2. Histone chromatin profiles of PRC2$^{+}$-CGI promoters in different normal tissues. (A)** H3K27ac and **(B)** H3K27me3 profiles of PRC2$^{+}$-CGI promoters. PRC2$^{+}$-CGI promoters are stratified into 5 groups based on gene expression in TCGA normal tissues of the corresponding cancer type.

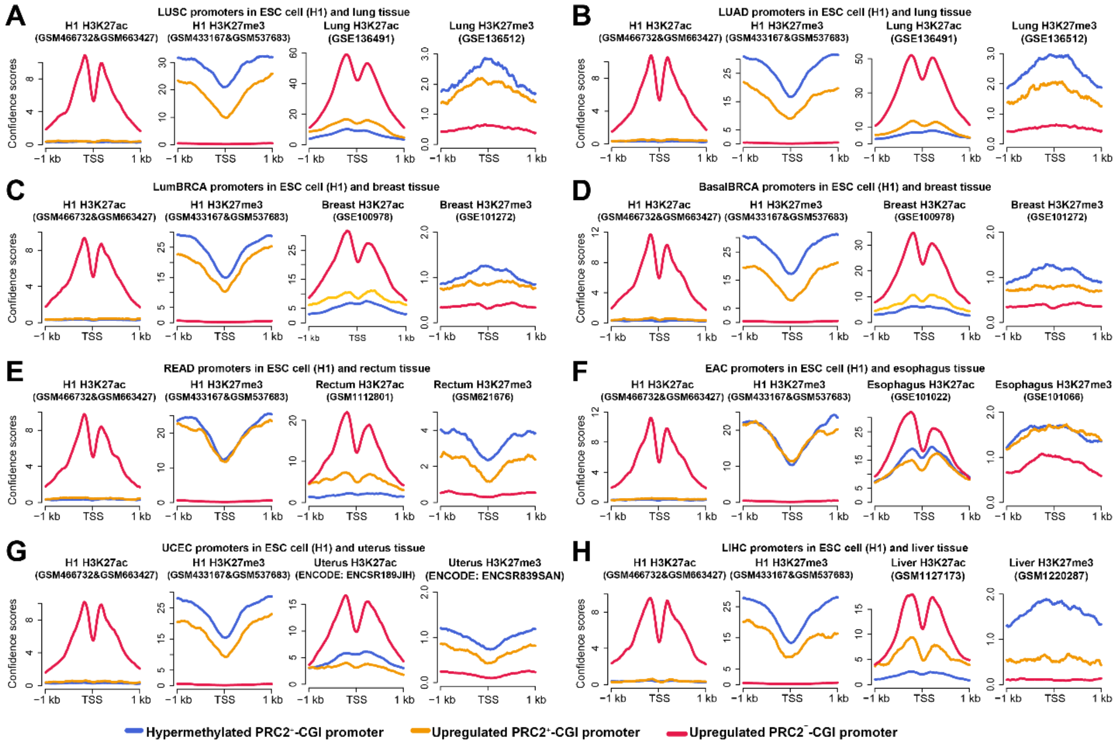


**Fig. S3. ESCs (H1 cells) and normal tissues harbor comparable levels of H3K27ac and H3K27me3 on the three classes of CGI promoters**

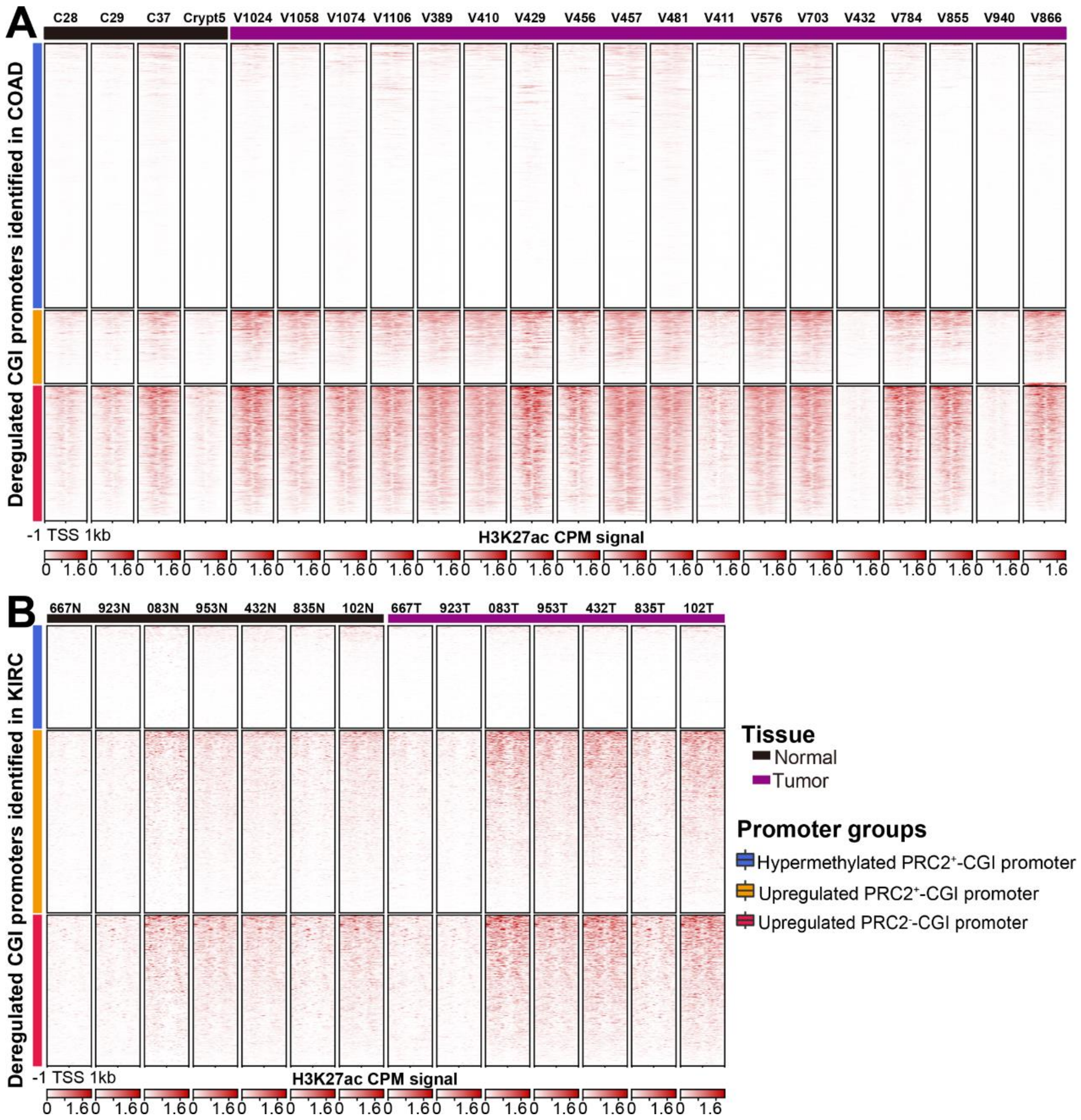


**Fig. S4. H3K27ac profiles of three different groups of CGI promoters in tumor and nonmalignant tissues from (A) COAD (GSE77737) and (B) KIRC (GSE86095) samples.**

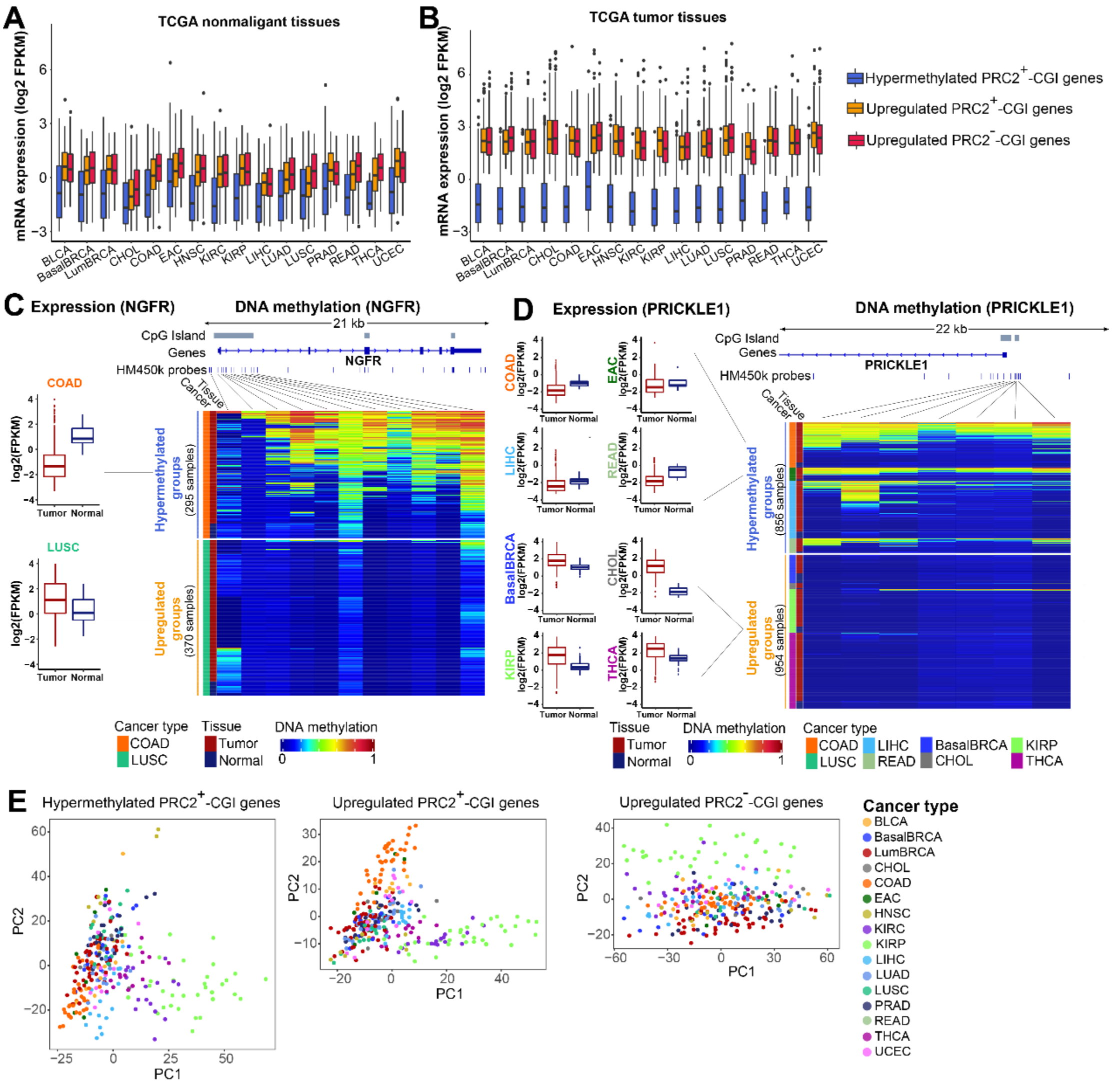


**Fig. S5. Upregulated PRC2$^{+}$-CGI genes have highest cancer-type-specificity and regulatory plasticity. (A-B)** Pan-cancer gene expression profiles of **(A)** nonmalignant and **(B)** tumor tissues across three CGI gene categories. **(C-D)** NFGR **(C)** and PRICKLE1 **(D)** are additional examples of plastic PRC2$^{+}$-CGI genes. **(E)** PCA analyses on the ATAC-Seq signal of different gene categories were performed across pan-cancer samples.

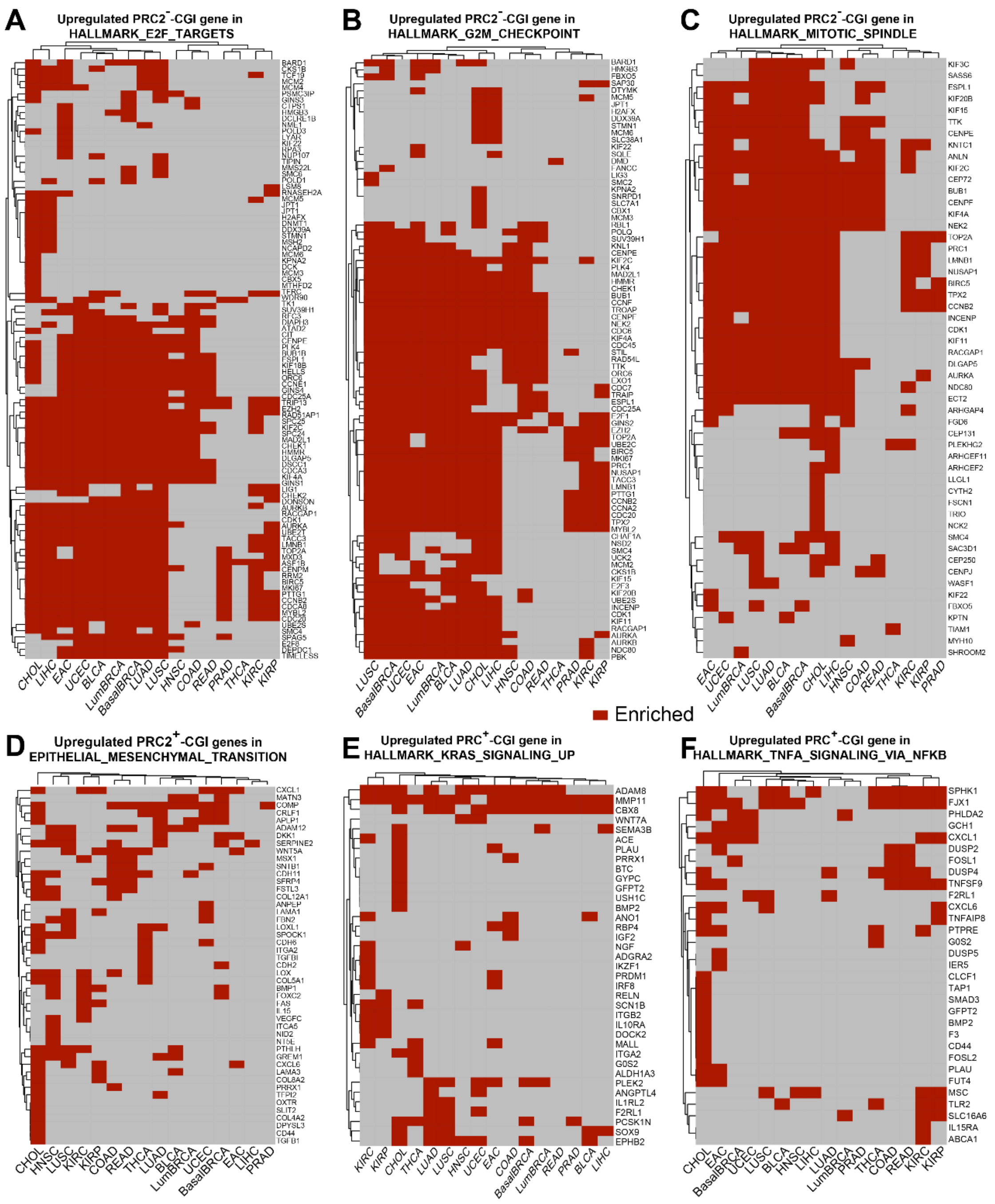


**Fig. S6. Heatmaps of the upregulated PRC2$^-$-CGI genes in E2F targets, G2M checkpoint as well as mitotic spindle pathways (upper) and PRC2$^+$-CGI genes in Epithelial mesenchymal transition, KRAS signaling up and TNFα signaling via NFKB pathways (lower) across cancer types.** Red bar indicates the enrichment of the gene in the pathway.

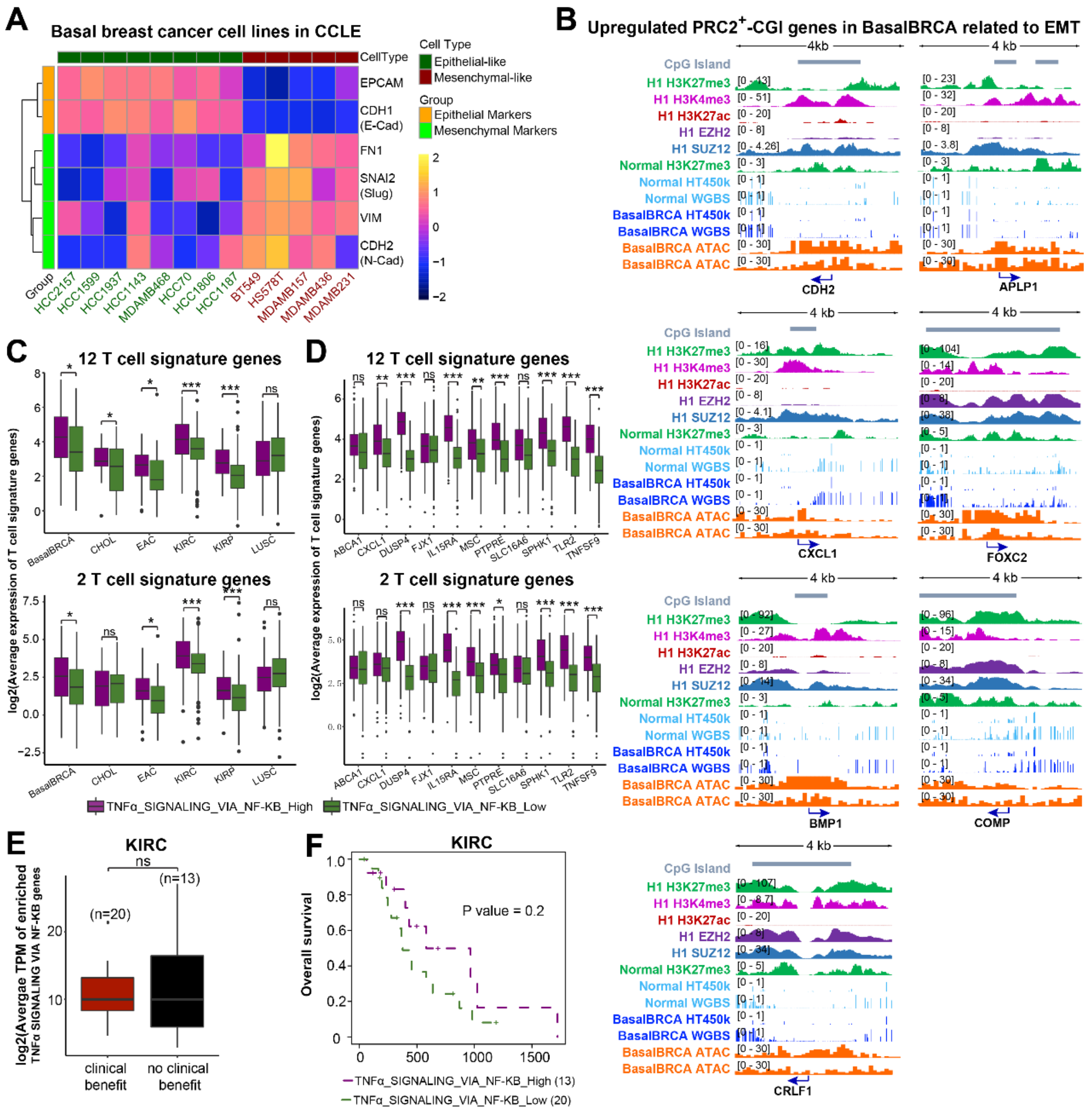


**Fig. S7. Upregulated PRC2$^{+}$- and PRC2$^{-}$-CGI genes control distinct sets of biological pathways in cancer. (A)** Known EMT markers were used to confirm the EMT state of epithelial-like and mesenchymal-like cells. **(B)** IGV plots show the remaining upregulated EMT-related genes in BasalBRCA. **(C-F)** Average expression of CD8+ T-cell signature genes in the six cancer types enriched in the "TNFα signaling via NF-KB" pathway, showing the top and bottom 20% of tumors based on either the average **(C)** or the individual gene-level **(D)** expression of upregulated PRC2$^{+}$-CGI genes in the "TNFα signaling via NF-KB" pathway. p values were determined by t-test. $p<0.001$, ***; $p<0.01$, **; $p<0.05$, *; $p>0.05$, ns. Expression datasets are obtained from Miao *et.al.* (35). **(E)** The average expression of upregulated PRC2$^{+}$-CGI genes in the "TNFα signaling via NF-KB" pathway in KIRC patients with differential response to immune checkpoint therapies. **(F)** Kaplan-Meier survival plot analyzing the average expression of upregulated PRC2$^{+}$-CGI genes in “TNFα signaling via NF-KB” pathway using the same cohort of KIRC patients.

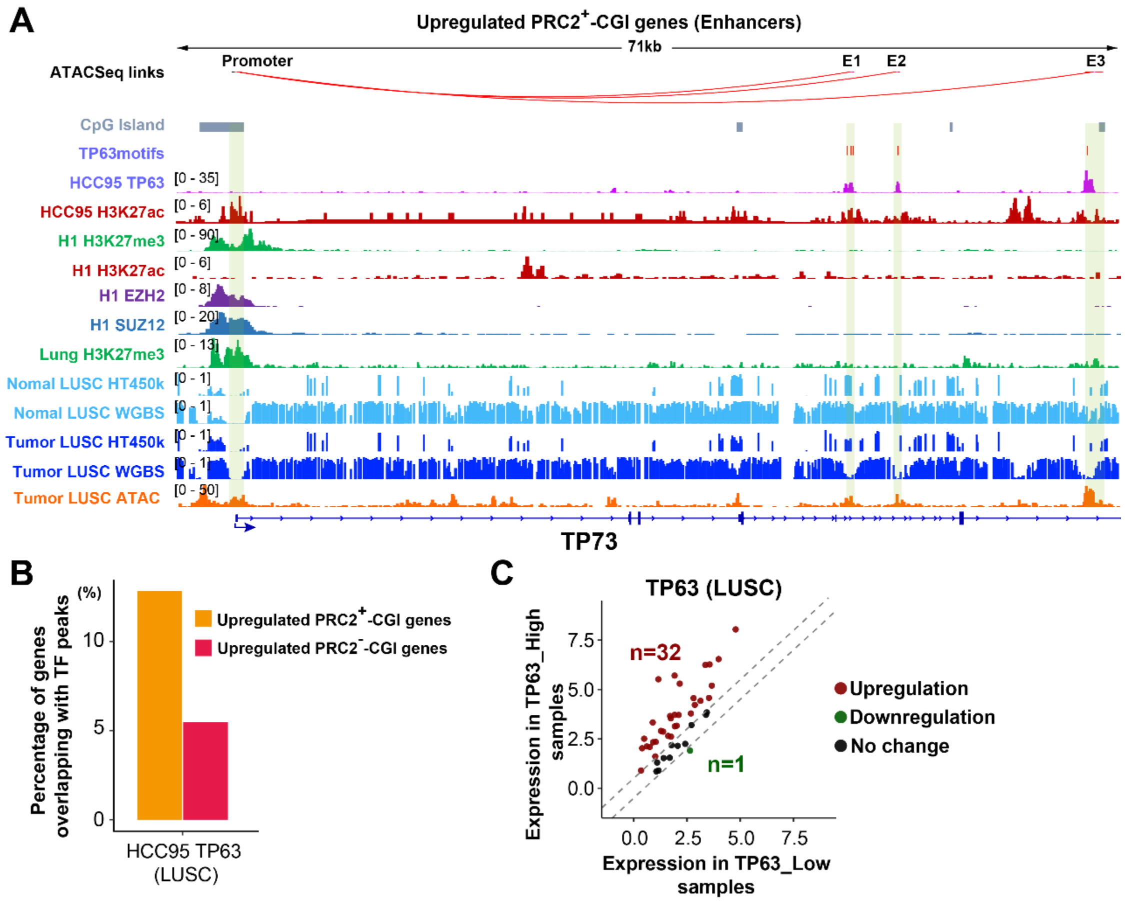


**Fig. S8. Upregulated PRC2$^{+}$-CGI genes are linked to distal enhancers targeted by specific transcription factor binding sites (TFBSs). (A)** TP73 enhancers (E1, E2, E3) predicted with TP63 motifs are occupied by TP63 in LUSC cells. TP63 ChIP-seq are from HCC95 (LUSC) cells (GSE66992). **(B)** TP63 ChIP-Seq peaks overlapping PRC2$^{+}$-CGI vs. PRC2$^{-}$-CGI enhancers, using the ChIP-Seq dataset above. **(C)** Expression differences between TCGA TP63-high and TP63-low LUSC tumors for the TP63 target genes having enhancers overlapped by TP63 in LUSC cells (from panel B). High and low tumors were those in the upper and lower quintile of TP63 expression.
